# Pulsed Third-Spin-Assisted Recoupling NMR for Obtaining Long-Range ^13^C-^13^C and ^15^N-^13^C Distance Restraints

**DOI:** 10.1101/2020.05.20.105221

**Authors:** Martin D. Gelenter, Aurelio J. Dregni, Mei Hong

**Affiliations:** Department of Chemistry, Massachusetts Institute of Technology, 170 Albany Street, Cambridge, MA 02139

## Abstract

We introduce a class of pulsed third-spin-assisted recoupling (P-TSAR) magic-angle-spinning (MAS) solid-state NMR techniques that achieve efficient polarization transfer over long distances to provide important restraints for structure determination. These experiments operate with the same principle as continuous-wave (CW) TSAR experiments, by utilizing second-order cross terms between strong ^1^H-^13^C and ^1^H-^15^N dipolar couplings to achieve ^13^C-^13^C and ^13^C-^15^N polarization transfer. However, in contrast to the CW-TSAR experiments, these pulsed P-TSAR experiments require much less radiofrequency (rf) energy and allow a much simpler routine for optimizing the rf field strength. We call the techniques PULSAR (PULSed proton Asissted Recoupling) for homonuclear spin pairs and ^PERSPIRATION^CP (Proton-Enhanced Rotor-echo Short Pulse IRradiATION Cross-Polarization) for heteronuclear spin pairs. We demonstrate these techniques on the model protein GB1, and found cross peaks for distances as long as 10 and 8 Å for ^13^C-^13^C and ^15^N-^13^C spin pairs, respectively. We also apply these methods to the amyloid fibrils formed by the peptide hormone glucagon, and show that long-range correlation peaks are readily observed to constrain intermolecular packing in this cross-β fibril. We provide an analytical model for the PULSAR and ^PERSPIRATION^CP experiments to explain the measured and simulated chemical shift dependence and pulse flip angle dependence of polarization transfer. These two techniques are useful for measuring long-range distance restraints to determine the three-dimensional structures of proteins and other biological macromolecules.

## Introduction

Magic-angle-spinning (MAS) solid-state nuclear magnetic resonance (SSNMR) spectroscopy produces high-resolution spectra by averaging orientation-dependent nuclear spin interactions such as chemical shift anisotropy (CSA), dipolar couplings, and quadrupolar couplings. MAS SSNMR is uniquely suited among spectroscopic techniques to provide atomic-resolution structural information of insoluble, non-crystalline biological macromolecules such as membrane proteins ^1–4^, amyloid fibrils ^5–13^, and cell walls ^14–17^. Once chemical shift assignment is complete, three-dimensional structure determination predominantly relies on inter-atomic distance restraints. These are typically measured through internuclear dipolar couplings. Under MAS, dipolar couplings are reintroduced using tailored radiofrequency (rf) pulse sequences or a careful choice of spinning frequency ^18^. Heteronuclear dipolar couplings and distances can be measured using the Rotational Echo DOuble Resonance (REDOR) technique ^19^ and the related Transferred Echo DOuble Resonance (TEDOR) ^20–21^, while homonuclear dipolar couplings and distances are typically measured qualitatively using spin diffusion techniques such as Proton-Driven Spin-Diffusion (PDSD) ^22^, Dipolar-Assisted Rotational Resonance (DARR) ^23–24^ and COmbined R2^ν^_n_ Driven spin diffusion (CORD) ^25^. Higher static magnetic field strengths are advantageous for obtaining higher spectral resolution and sensitivity. However, at higher fields the larger isotropic chemical shift differences weaken the spin diffusion mechanism. Moreover the larger CSA at high fields also calls for faster MAS, which further suppresses the dipolar couplings that drive homonuclear polarization transfer. In addition, nuclei with low gyromagnetic ratios (γ) have weak dipolar couplings, which make it difficult to measure the long distances that are particularly useful for constraining 3D structures. Therefore, it is important to develop improved pulse sequences for high-field fast MAS conditions to achieve efficient long-range polarization transfer ^26^.

Third-Spin-Assisted Recoupling (TSAR) experiments are a class of techniques that transfer polarization between low-γ nuclei A and B ^27^. This transfer occurs via cross terms in the second-order average Hamiltonian between the H–A and H–B dipolar couplings, which are generally much stronger than the direct A-B dipolar coupling. In continuous-wave TSAR (CW-TSAR) experiments, the homonuclear variant is known as Proton-Asisted Recoupling (PAR) ^28^ whereas the heteronuclear variant is called Proton-Asissted Insensitive Nuclei Cross Polarization (^PAIN^CP)^29–30^. The CW irradiation in these experiments requires carefully optimized rf field strengths on both the ^1^H channel and the heteronuclear channels. This is usually achieved with an extensive multidimensional search of the rf field strengths, without which the polarization transfer efficiency can easily fall below the theoretical maximum ^31^. Moreover, mixing times of 10–30 ms are usually required to observe long-range ^13^C-^13^C and ^15^N-^13^C correlation peaks. The extended high-power CW irradiation on two to three channels can damage the NMR probe and heat-sensitive biological samples. These two practical drawbacks have limited the use of the PAR and ^PAIN^CP experiments, despite their theoretical ability for long-range spin polarization transfer, compared to the simpler spin diffusion experiments.

Here we introduce pulsed TSAR (P-TSAR) techniques that achieve efficient polarizarion transfer over long distances using low rf energy and simple optimization routines. These P-TSAR experiments rely on a pulsed spin-lock^32^, in contrast to CW-TSAR experiments that utilize a CW spin-lock. We demonstrate these techniques on a model protein, the β1 immunoglobulin binding domain of protein G (GB1), and apply them to the amyloid fibrils formed by the peptide hormone glucagon ^12^. We then show by numerical simulations, average Hamiltonian Theory ^33^ and an analytical model that these P-TSAR experiments have the same essential spin dynamics as CW-TSAR: they have vanishing first-order average Hamiltonians ^33^ and rely on trilinear terms of the kind 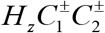 and *H_z_C*^±^*N*^±^ in the second-order average Hamiltonian to mediate polarization transfer. We call the homonuclear experiment PULSed proton-Asissted Recoupling (PULSAR) **(Fig. 1a)** and the heteronuclear experiment Proton-Enhanced Rotor-echo Short Pulse IRradiATION (^PERSPIRATION^CP). The latter was introduced recently by us ^34^ but is now significantly improved by altering the timing of the spin-lock pulses to better refocus the magnetization at the end of each rotor period (**Fig. 1c**), thus obviating the need for z-filters. Most importantly, P-TSAR experiments use only 10-20% of the rf duty cycle of their CW-TSAR analogs on the low-frequency channels and are robust against variations in the heteronuclear rf powers.

**Figure 1.**
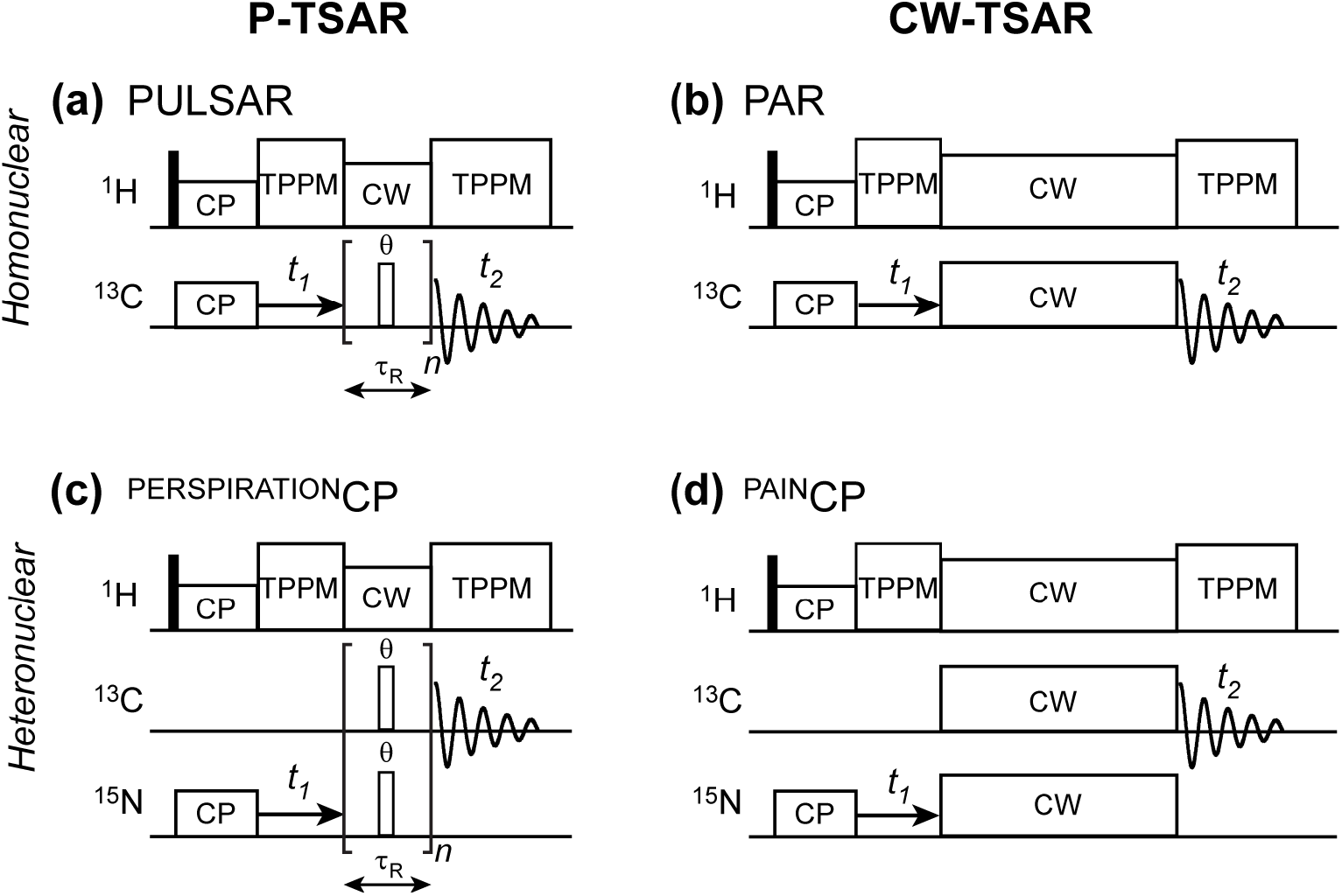
P-TSAR pulse sequences developed in this study (**a, c**), compared with the previously published CW-TSAR pulse sequences (**b, d**). **(a)** 2D ^13^C-^13^C PULSAR. **(b)** 2D ^13^C-^13^C PAR ^28^. **(c)** 2D ^15^N-^13^C ^PERSPIRATION^CP with the spin-lock pulses in the middle of the rotor period. **(d)** 2D ^15^N-^13^C ^PAIN^CP ^28^.

## Materials and Methods

SSNMR experiments were conducted on Bruker Avance III HD 600 MHz (14.1 T) and Avance II 800 MHz (18.8 T) spectrometers equipped with triple-resonance ^1^H/^13^C/^15^N 3.2 mm MAS probes. All experiments were carried out under 20 kHz MAS. Microcrystalline GB1 samples were expressed and purified as previously described ^35–36^. Glucagon peptide was purchased from Biopeptek Pharmaceuticals (Malvern, PA) and fibrillized in pH 2 solution at 21°C for 7 days at a concentration of 8 mg/ml ^37^.

PULSAR and ^PERSPIRATION^CP experiments (**Appendix S4,5**) were conducted at a set temperature of 278 K for GB1 and 258-263 K for the anhydrous tripeptide formyl-Met-Leu-Phe (f-MLF) and for hydrated glucagon fibrils. At 20 kHz MAS, the actual sample temperature is estimated to be ~15 K higher than the set temperature due to frictional heating ^38^. Two-Pulse Phase Modulation (TPPM) ^1^H decoupling ^39^ was applied at an rf field of 71-83 kHz during ^13^C detection. For 2D ^13^C-^13^C PULSAR experiments, the direct acquisition times were 21.5 ms and 16.1 ms for the 600 and 800 MHz spectra, respectively, whereas the indirect dimension ^13^C evolution times were 9.9 ms and 7.4 ms, respectively. For the 2D ^15^N-^13^C ^PERSPIRATION^CP experiments, the ^13^C acquisition time was 11.9 ms and the ^15^N evolution time was 10.2 ms. A pseudo-3D ^PERSPIRATION^CP experiment was conducted on the 800 MHz spectrometer in which one indirect dimension varied the ^13^C rf carrier from 0-200 ppm in 2.5 ppm increments, and the other indirect dimension varied the spin-lock pulse flip angles for both ^13^C and ^15^N from 0 to 360° in 15° increments. For the spin-lock pulses, the ^13^C and ^15^N rf field strengths were 50 kHz and 38.8 kHz, respectively, while the ^1^H CW rf irradiation was 56–59 kHz during PULSAR and ^PERSPIRATION^CP polarization transfer.

All 2D spectra were processed in TopSpin (Bruker BioSpin) and assigned in Sparky ^40^. The distance upper limits were set to 10 Å for the ^13^C-^13^C cross peaks in the PULSAR spectra and 8 Å for the ^13^C-^15^N cross peaks in the ^PERSPIRATION^CP spectra. We calculated the GB1 structure using CYANA 2.1 ^41^. (ϕ, ψ) torsion angles were predicted from backbone chemical shifts using the TALOS-N software ^42^. The angular uncertainty was set to twice the uncertainty given by TALOS-N. The structure calculation consisted of a total of 100 independent runs with 70,000 torsion angle steps each. The ten structures with the lowest target function were included in the final ensemble. Numerical simulations of the PULSAR and ^PERSPIRATION^CP polarization transfer dynamics were conducted using SpinEvolution ^43^. Analytical calculations and data analysis were performed using home-written MATLAB and Python scripts, which can be found in the Supporting Information. The magnetization trajectories and Bloch spheres were generated in MATLAB.

## Results

### Demonstration of PULSAR and ^PERSPIRATION^CP experiments on a microcrystalline model protein

**Fig. 1** shows the pulse sequences for PULSAR and ^PERSPIRATION^CP and their CW-TSAR counterparts. Both P-TSAR experiments use pulses with a flip angle θ at the center of each rotor period on the low-frequency channels. CW decoupling occurs on ^1^H during the mixing time. As in PAR and PAINCP, the ^1^H decoupling field strength is chosen to avoid the rotary resonance condition, ω_1_=ω_r_, and the CP matching condition with the heteronuclear channels. We conducted all experiments under 20 kHz MAS and used rf field strengths of 35-50 kHz for the θ-pulses.

We first demonstrate the PULSAR and ^PERSPIRATION^CP experiments on the model protein GB1 on a 600 MHz spectrometer **(Fig. 2)**. We placed the ^13^C carrier frequency at 43 ppm, in the middle of the aliphatic region, and the ^15^N carrier frequency at 122 ppm, in the middle of the amide region. We varied the spin-lock pulse flip angle from 90° to 165°, and found 150° pulses to give maximum polarization transfer in a broadband fashion. The aliphatic ^13^C chemical shift range for which intensity is observed is about 9 kHz (60 ppm). Intensity is also observed in the aromatic and carbonyl regions of the spectrum. At 600 MHz, the aliphatic-carbonyl chemical shift difference is ~20 kHz (130 ppm). Smaller flip angles reduced the chemical shift range of the polarization transfer, whereas larger flip angles decreased the total transferred magnetization. The optimal ^1^H rf field strength during the ^13^C and ^15^N P-TSAR period is slightly less than 3*ω_r_* for both PULSAR and PERSPIRATIONCP experiments, consistent with numerical simulations **(Fig. S2)**. The 2D ^13^C-^13^C PULSAR spectrum shows many long-range correlations in every region of the 2D spectrum. These include aliphatic-aliphatic, aliphatic-aromatic, aliphatic-carbonyl, aromatic-aromatic correlations (**Fig. 2a**). Some of the cross peaks correspond to distances of 8 Å or more, such as T25-Y3 (**Fig. 2b**). The 2D ^15^N-^13^C ^PERSPIRATION^CP spectrum similarly showed many long-range correlations (**Fig. 2c**) such as the V54-L7, which has a distance of 6 Å (**Fig. 2d**).

**Figure 2.**
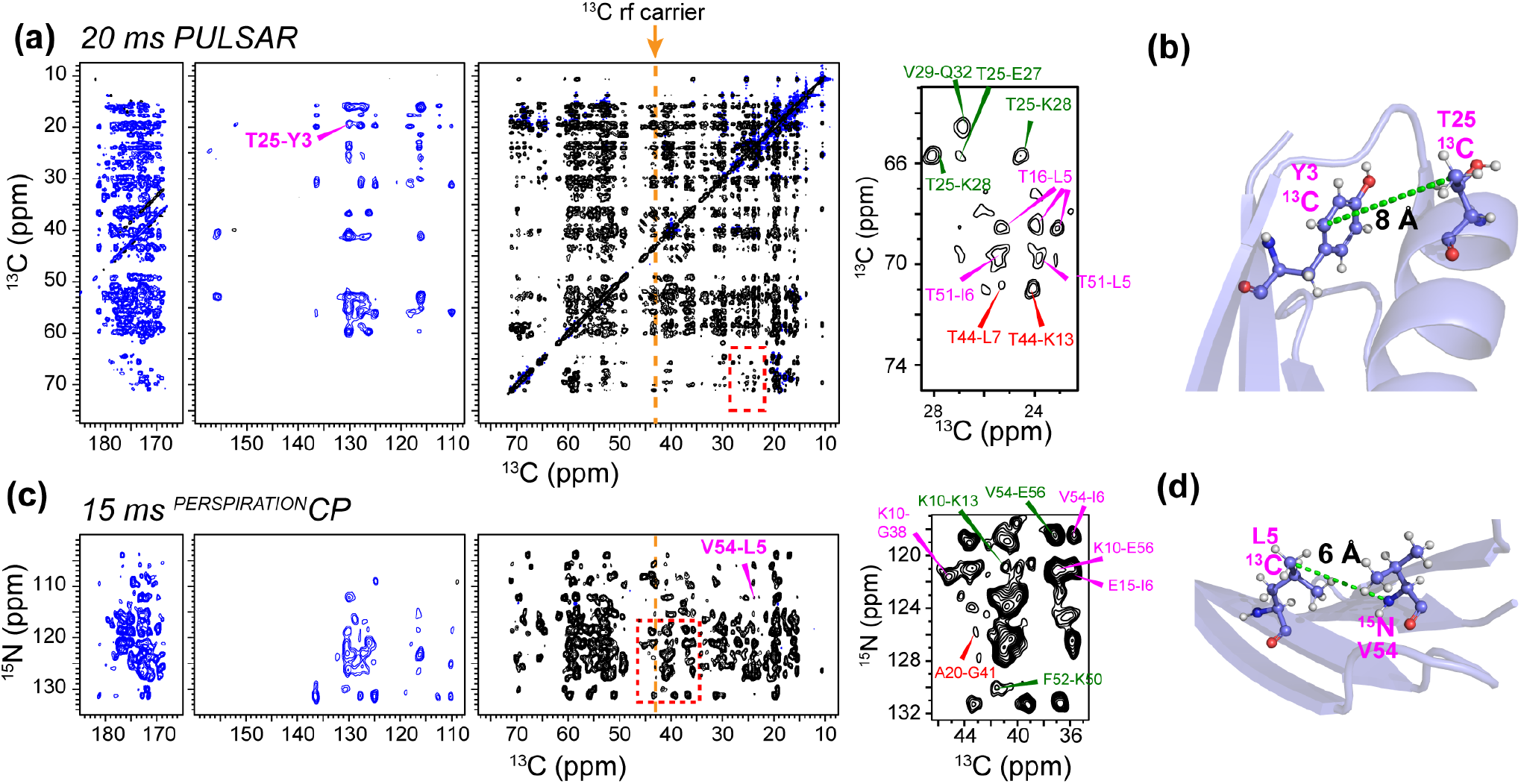
2D PULSAR and ^PERSPIRATION^CP spectra of GB1, measured on the 600 MHz spectrometer. **(a)** 2D ^13^C-^13^C PULSAR spectrum measured with 20 ms mixing. The ^13^C rf carrier frequency was set at 42 ppm (orange arrow). Positive intensities are shown in black and negative intensities are shown in blue. The inset on the right illustrates representative medium-range (green), long-range (pink), and intermolecular (red) correlation peaks. **(b)** GB1 structure, showing one of the long-range contacts, between T25 and Y3, whose cross peak is annotated in (**a**). **(c)** 2D ^15^N-^13^C ^PERSPIRATION^CP spectrum measured with a 15 ms mixing time and 150° recoupling pulses. Zoomed-in area shows representative observed medium-range (green), long-range (pink), and intermolecular (red) correlation peaks. **(d)** Structure of GB1 showing one of the ^15^N-^13^C contacts, between V54 and L7, whose cross peak is annotated in (**c**).

Because of the enhanced sensitivity and resolution provided by higher magnetic fields, we also conducted PULSAR experiments on an 800 MHz spectrometer, where the aliphatic chemical shifts span a range of 12 kHz and the aliphatic-carbonyl chemical shift separation increases to 26 kHz. To achieve the same offset-dependent polarization transfer as the 600 MHz spectra, one would need to spin the samples at 27 kHz. Since this is not feasible using 3.2 mm MAS rotors and nitrogen gas for spinning, we kept the MAS rate at 20 kHz and varied the pulse flip angle to optimize polarization transfer in the spectral region of interest. With 150° pulses, ^13^C magnetization from the aliphatic carbons gave positive transfer to other aliphatic carbons, negative transfer to aromatic carbons (110-156 ppm), and positive transfer to carbonyl peaks (172-182 ppm) (**Fig. 3a**). With 210° pulses, the transfer profile has a null in the aromatic region and negative intensities in the carbonyl region (**Fig. 3b**). Similarly long distances were observed as spectra measured at 600 MHz.

**Figure 3.**
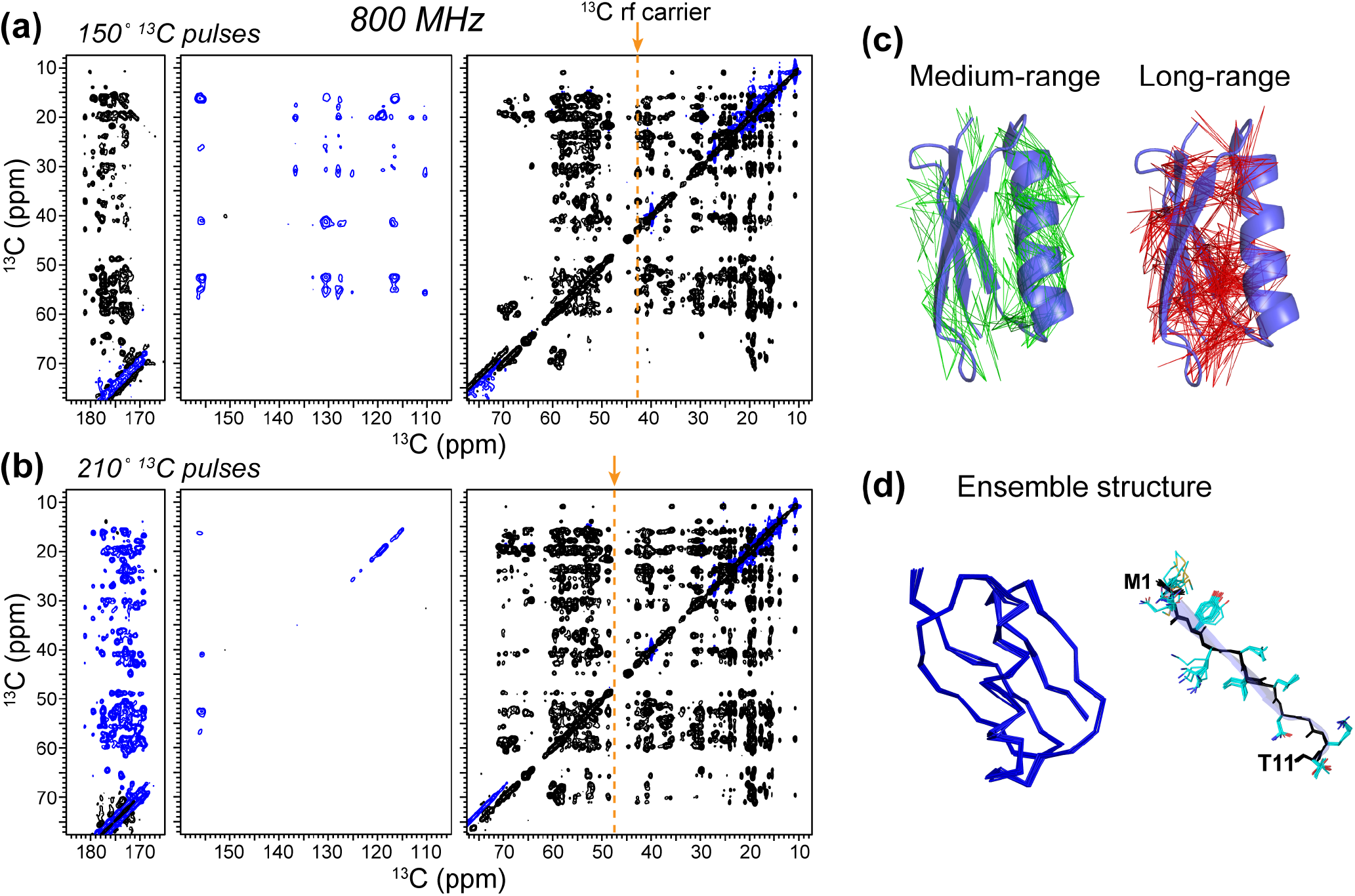
2D ^13^C-^13^C PULSAR spectra of GB1 measured at 800 MHz to show the offset dependence of PULSAR. **(a)** Spectra measured using 150° pulsed spin-lock pulses. (**b)** Spectra measured using 210° pulsed spin-lock pulses. The rf fields were 50 kHz for ^13^C and 56 kHz for ^1^H and the mixing time was 20 ms. **(c)** 112 medium-range (green) and 176 long-range (red) correlations in GB1 obtained from the two PULSAR and ^PERSPIRATION^CP spectra. **(d)** Lowest-energy ensemble of GB1 structures calculated based on the distance constraints from PULSAR and ^PERSPIRATION^CP spectra. The backbone ensemble (left) shows that P-TSAR distance restraints combined with (φ,ψ) dihedral angles is sufficient to restrain the 3D structure of the protein. The M1-T11 φψ-strand is shown on the right to illustrate the sidechain conformations constrained by the measured long-range correlations.

These 2D ^13^C-^13^C and ^15^N-^13^C correlation spectra gave rise to a total of 100 sequential, 112 medium-range (2 < |*i-j*| < 5), and 176 long-range (|*i-j*| > 5) distance restraints (**Fig. 3c**). Combining these restraints with (ϕ, ψ) torsion angles obtained from backbone chemical shifts ^42^, we calculated the GB1 structure. We set the upper distance limits to 10 Å for PULSAR cross peaks and 8 Å for ^PERSPIRATION^CP cross peaks. These upper limits were estimated by finding cross peaks corresponding to the longest distance in the high-resolution NMR structure of GB1^44^ and ensuring that this upper limit does not cause restraint violations in the structure calculation. The medium-range restraints are particularly useful for determining the secondary structure, since *i* to *i* + 2 cross peaks are expected for β-strands while *i* to *i +* 3 or 4 cross peaks are expected for θ-helices (**Fig. 3c, *left***). The long-range restraints helped to define both the backbone fold and the sidechain conformations (**Fig. 3c, *right***). The ten lowest-energy structures display a backbone RMSD of 0.4 ± 0.1 Å and a heavy-atom RMSD of 1.3 ± 0.2 Å (**Fig. 3d**). Therefore, these ^13^C-^13^C PULSAR and ^15^N-^13^C ^PERSPIRATION^CP restraints are sufficient to solve the high-resolution structure of GB1.

### Application of P-TSAR to glucagon amyloid fibrils

To test the utility of the ^13^C-^13^C PULSAR and ^15^N-^13^C ^PERSPIRATION^CP experiments for structure determination of non-crystalline proteins, we applied them to the amyloid fibrils formed by the peptide hormone glucagon ^12^. Glucagon is used as a compound to treat severe hypoglycemia. At the high concentrations required for intravenous administration, it fibrillizesrapidly ^45^ into cross-β structures. This fibrillization tendency has limited glucagon’s formulation to a dry powder, to be mixed in acidic solution immediately prior to injection. Understanding the atomic structure of glucagon fibrils is thus important for designing glucagon analogs that resist fibril formation. We measured the 2D PULSAR and ^PERSPIRATION^CP spectra of a glucagon fibril containing nine ^13^C, ^15^N-labeled residues (**Fig. 4a**). The 2D ^13^C-^13^C PULSAR spectrum was measured at 800 MHz using a 12.5 ms mixing time (**Fig. 4b**) while the 2D ^15^N-^13^C ^PERSPIRATION^CP spectrum was measured at 600 MHz using a 15 ms mixing time (**Fig. 4c**). Many medium-range intramolecular cross peaks were observed, such as T5Cβ-Q3Cβ and Q24Cγ-L26Cδ1 (**Fig. 4b**). This is consistent with the location of *i* and *i + 2* sidechains on the same side of the β-sheet plane. We also detected long-range correlations such as T5Cθ-M27Cθ and T5Cθ-L26Cδ in the PULSAR spectrum and G4N-N28Cθ and W25N-M27Cεcross peaks in the ^15^N-^13^C correlation spectra. These long-range correlations can only be assigned to intermolecular contacts, based on additional experiments of isotopically diluted samples ^12^. Therefore, the long-range correlations define the antiparallel packing of β-strands. Intra-residue backbone ^15^N to sidechain ^13^C correlations are also observed for bulky sidechains such as Q24N-W25Cζ_3_. These restraints combine with intermolecular correlations involving the same W25 sidechain to determine the rotameric structure of the bulky sidechain (**Fig. 4d, e**).

**Figure 4.**
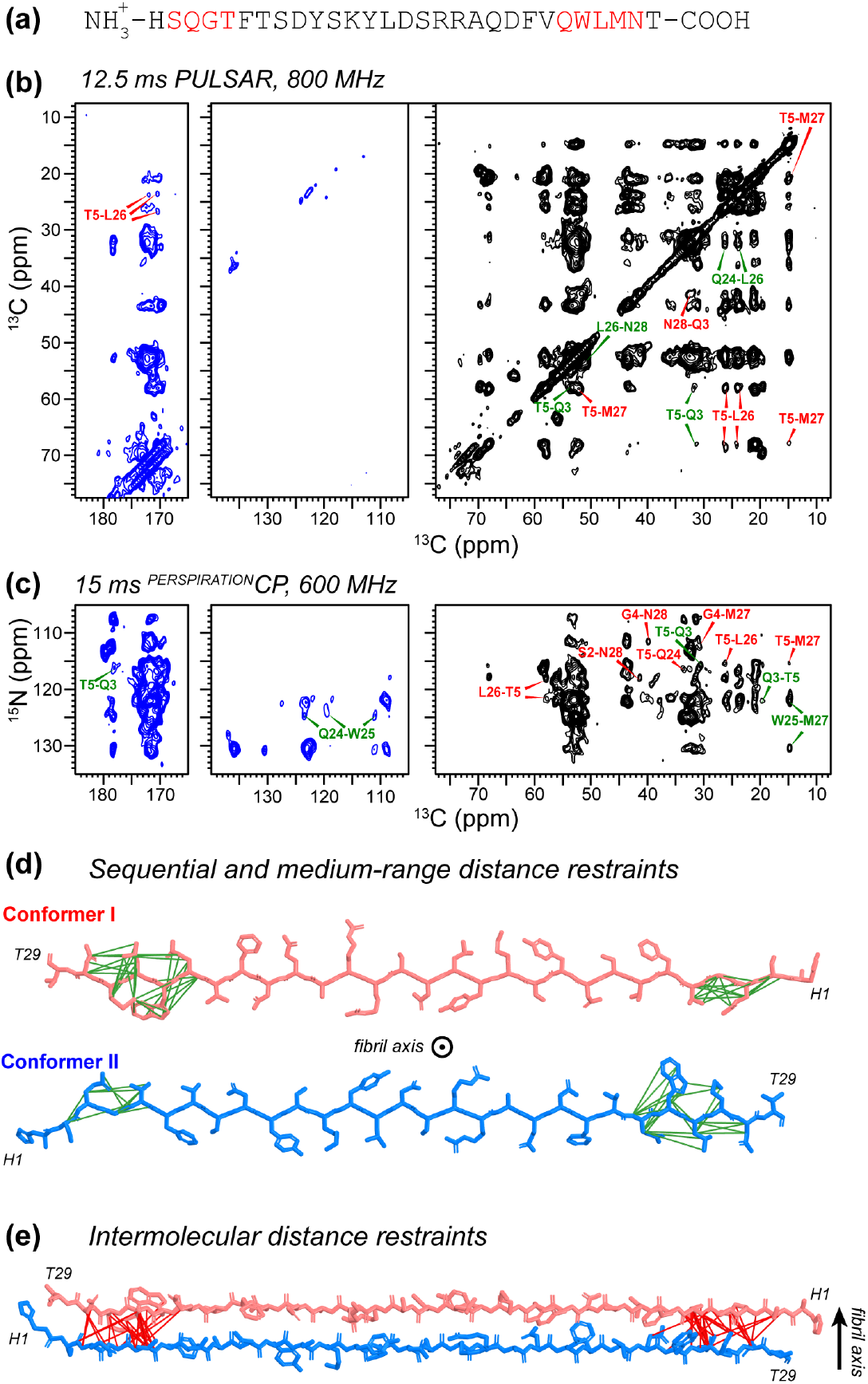
2D PULSAR and ^PERSPIRATION^CP experiments on glucagon amyloid fibrils for obtaining distance restraints. **(a)** Amino acid sequence of the glucagon peptide. The ^13^C, ^15^N-labeled residues are shown in red. **(b)** 12.5 ms 2D ^13^C PULSAR spectrum, measured on the 800 MHz spectrometer using 150° ^13^C recoupling pulses. **(c)** 15 ms 2D ^15^N-^13^C ^PERSPIRATION^CP spectrum measured on the 600 MHz spectrometer using 150° ^13^C and ^15^N recoupling pulses. **(d)** Sequential and medium-range intramolecular correlations found in the PULSAR and ^PERSPIRATION^CP spectra. These correlations help to define the rotomeric conformation of bulky sidechains such as W25. **(e)** Intermolecular correlations define antiparallel hydrogen bonding of the β-strands along the fibril axis.

### Numerical simulations of CW-TSAR and P-TSAR experiments

We conducted numerical simulations to understand the chemical shift offset and rf field dependences of the P-TSAR experiments compared to CW-TSAR. A 7-spin system was used for PULSAR and PAR while an 8-spin system was used for ^PERSPIRATION^CP and ^PAIN^CP. These spin systems were taken from amino acid Met to represent the backbone and sidechain structure of a typical amino acid residue (**Fig. 5a, d**). For PULSAR and ^PERSPIRATION^CP experiments, we simulated the polarization transfer as a function of ^13^C isotropic chemical shift offset of the observed (sink) spins and as a function of the pulse flip angle. For PAR and ^PAIN^CP experiments, we simulated the transfer efficiency as a function of the rf field strength. Representative simulation scripts are given in Supporting Information **Appendix S1**.

**Figure 5.**
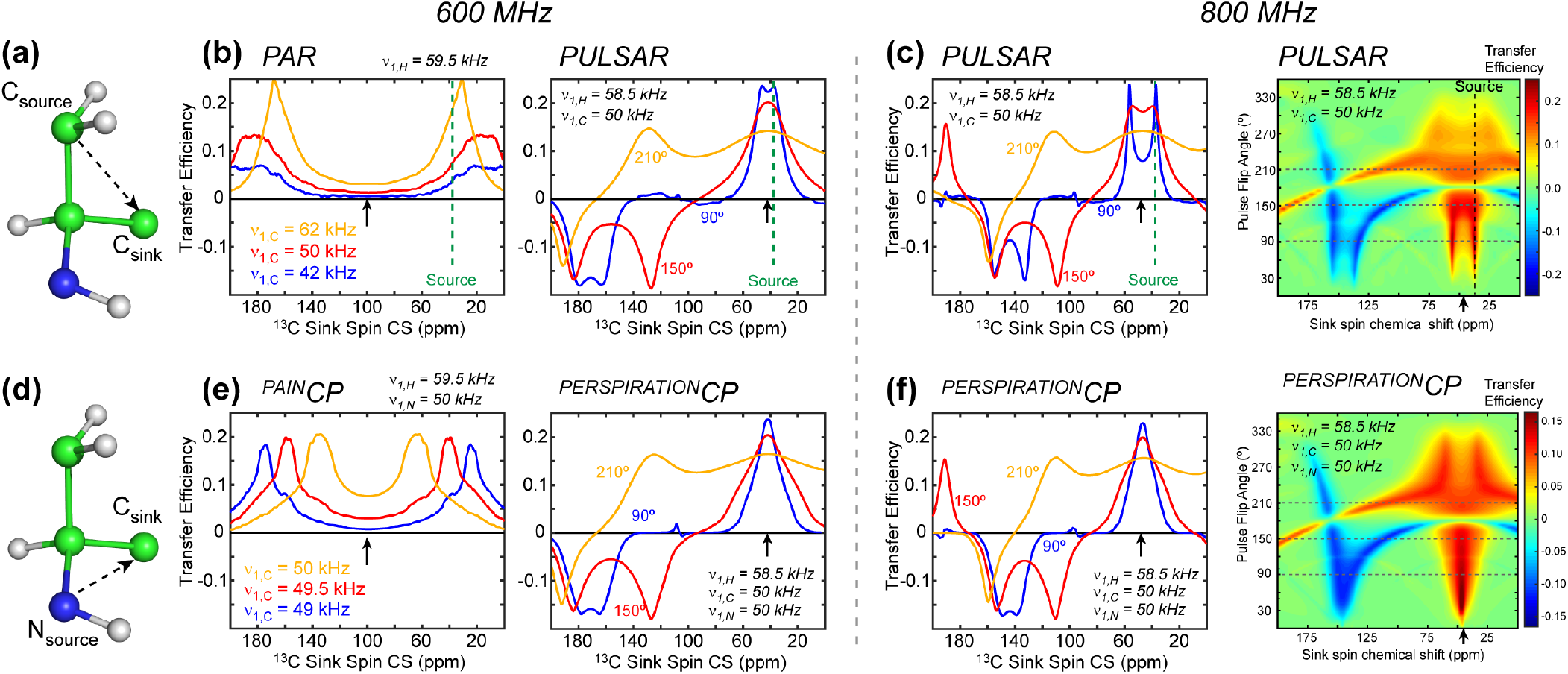
Numerical simulations of the chemical shift dependence of TSAR experiments. A mixing time of 20 ms was used in all simulations. **(a)** 7-spin system for simulating the PAR and PULSAR ^13^C-^13^C polarization transfer. The ^15^N spin is excluded in these simulations. **(b)** PAR and PULSAR efficiencies at 600 MHz as a function of the observed ^13^C chemical shifts. The ^13^C rf carrier frequency is denoted with a black arrow while the ^13^C source spin chemical shift is denoted with a green dashed line. **(c)** PULSAR transfer efficiency 800 MHz. (right) 2D plot of the PULSAR transfer efficiency as a function of sink spin chemical shift and pulsed spin-lock flip angle. **(d)** 8-spin system used for simulating ^PAIN^CP and ^PERSPIRATION^CP ^15^N-^13^C polarization transfer. **(e)**^PAIN^CP and ^PERSPIRATION^CP efficiencies at 600 MHz as a function of the observed ^13^C chemical shifts. **(f)**^PERSPIRATION^CP transfer efficiencies at 800 MHz. (right) 2D plot of the ^PERSPIRATION^CP transfer as a function of sink-spin chemical shift and pulse flip angle.

PAR transfer is optimized when the source and sink spin have similar ^13^C chemical shift offsets. With the ^13^C rf carrier frequency at 100 ppm, the PAR polarization transfer from a CH_2_ carbon at 40 ppm is efficient for aliphatic signals close to the source ^13^C chemical shift and for carbonyl carbons, but the transfer is minimal at about 100 ppm (**Fig. 5b, *left***). Over the range of ^13^C rf field strength from 62 to 42 kHz, stronger rf fields give higher transfer efficiencies. ^15^N-^13^C ^PAIN^CP exhibits a similar offset dependence (**Fig. 5e, *left***), but the transfer efficiency is much more sensitive to the ^13^C rf field strength: significantly higher transfer efficiency is found at a ^13^C rf field of 50 kHz than at 49 kHz, while ^1^H and ^15^N rf fields are held constant at 59.5 kHz and 50 kHz, respectively. The same modest ^13^C rf field change also changed the offset at which maximum transfer occurs, from 60 ppm to 30 ppm. Thus, ^PAIN^CP is extremely sensitive to both the rf field strength and the chemical shift offset.

Compared to these CW-TSAR polarization transfer profiles, the P-TSAR experiments give more broadband transfer and lower sensitivity to rf field strength. Maximum PULSAR transfer is also found when the source and sink spin have similar chemical shift offsets (**Fig. 5b, *right*** & **Fig. 5c**). However, unlike PAR, there are other matching conditions that also can accomplish polarization transfer. Prudent choice of pulse flip-angle and carrier frequency can allow an experiment to have a much wider range of chemical shift offsets with efficient transfer. At 600 MHz, when the ^13^C carrier is set to 38 ppm, the optimal ^13^C flip angle is ~150°, at which efficient transfer is found for all three regions of the ^13^C spectra: aliphatic, aromatic, and carbonyl (**Fig. 5b, *right***). When the flip angle increases to 210°, the aromatic signals becomes positive while the carbonyl intensities vanish. At 800 MHz, the highest polarization transfer is found at a flip angle of 210°, where positive transfer is observed from 0-130 ppm and negative transfer is observed for carbonyl carbons (**Fig. 5c**). The ^15^N-^13^C ^PERSPIRATION^CP transfer profiles (**Fig. 5e, *right*** & **Fig. 5f**) are similar to the PULSAR transfer profiles.

### Average Hamiltonian analysis of P-TSAR

To further understand the mechanism of polarization transfer in PULSAR and ^PERSPIRATION^CP, we analyzed their average Hamiltonians ^33^ using a similar approach to the CW-TSAR treatment ^28, 30^. For clarity, we first analyze ^PERSPIRATION^CP; explanations on how the analysis changes for PULSAR are given in the main text and Supporting Information. We consider a three-spin system consisting of one ^1^H, one ^15^N and one ^13^C. The ^1^H spin is subjected to CW rf irradiation while δ-function pulses with flip angles θ_*c*_ amd θ_*N*_ are applied at the center of each rotor period to ^13^C and ^15^N, respectively (**Fig. 1a,c**). The Zeeman-truncated internal Hamiltonian for ^PERSPIRATION^CP is

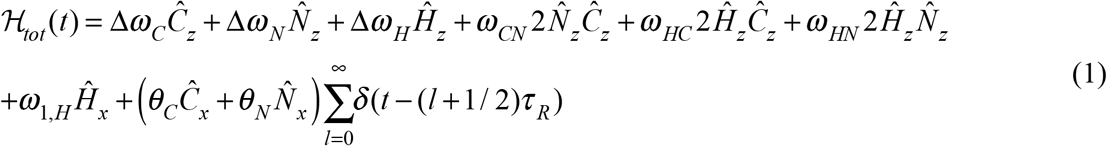

Here Δ*ω_C_* and Δ*ω_N_* are the sum of the isotropic chemical shift offset and chemical shift anisotropy (CSA) of ^13^C and ^15^N, ω_CN_ ω_HC_ω_HN_ are heteronuclear dipolar couplings, and ω_1,*H*_ is the ^1^H CW rf field strength. The Dirac δ(*t* − (*l* + 1 / 2)τ_*R*_) specifies the ^13^C and ^15^N pulses in the middle of each rotor period, *τ_R_*. MAS introduces time-dependence in the CSA and dipolar couplings, which is not explicitly shown in Eq. (1).

We transform this Hamiltonian into the interaction frame defined by the rf pulses for ^1^H and the frame defined by both rf pulses and isotropic chemical shift offset for ^13^C and ^15^N. We neglect ^1^H isotropic chemical shift because CW irradiation truncates chemical shift evolution. In contrast, the heteronuclear spins undergo chemical shift evolution between pulses. This interaction-frame transformation can be written as:

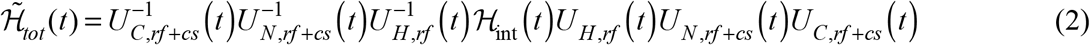

Here 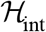 represents all the terms in Eq. (1) except for the rf pulses and the ^15^N and ^13^C isotropic chemical shift. The interaction-frame transformations in Eq. (2) each only affect one spin. As such, we can separately transform each one-spin operator in the Hamiltonian, and multiply the results together to obtain the two-spin product operators. We thus calculate the effect of the interaction frame transformation on the product operators 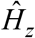, 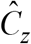, and 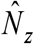.

For the ^1^H spin we consider the interaction frame of the ^1^H CW rf irradiation. Similar to the treatment of PAR^28^, we rotate 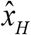 to 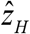 so that the spin-lock direction is 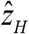. We convert product operators to spherical tensor operators:

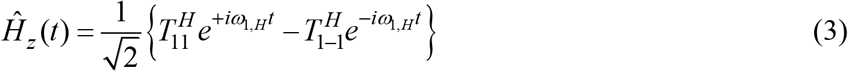

The propagator for the ^13^C rf pulse and isotropic shift is:

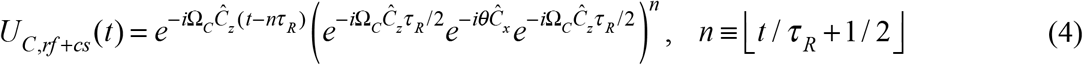

where *n* is the number of elapsed pulses defined by the floor function. Since any series of successive rotations is equivalent to a single rotation about a new axis, we define a pulsed spin-lock axis 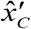 in the *x*, *z* plane, at an angle ψ_C_ from the *x*-axis towards the *z*-axis, and call the net rotation about this axis in each rotor period *φ_C_*. Each ^13^C spin in the system has a distinct isotropic chemical shift, and thus also has a distinct pulsed spin-lock direction 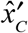 and net rotation per rotor period *φ_C_*. The propagator in Eq. (4) can then be simplified as:

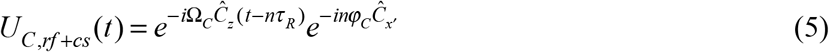

We next consider the interaction-frame transformation of 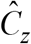:

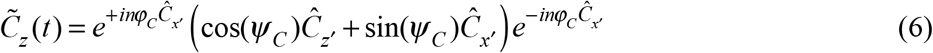

Similar to ^1^H, we rotate 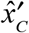 to 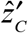 make the pulsed spin-lock direction 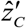, and convert product operators to spherical tensor operators:

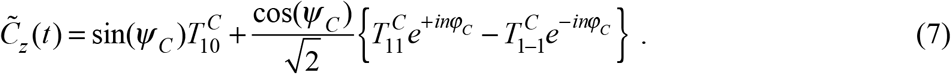

Here, for clarity, we have dropped the primes for the rotated axes on the spherical tensor operators. The interaction-frame ^15^N operators are identical to Eq. (4)-(7) except with all C replaced by N: the ^15^N spin has its own pulsed spin-lock direction 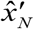, and the interaction-frame 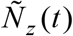 is given by an analogous equation to Eq. (7). The interaction-frame Hamiltonian (**Eq. (2)**) can thus be calculated by replacing the spin operators in Eq. (1) with their interaction-frame representations, Eq. (3) and (7):

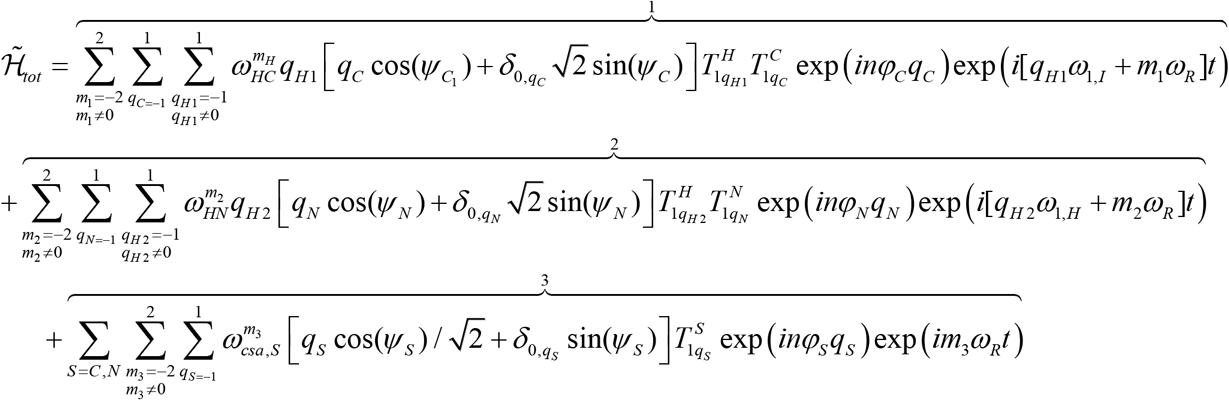

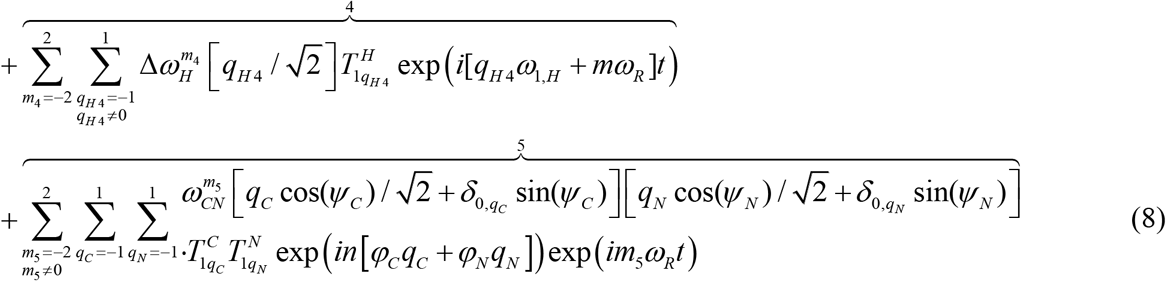

In Eq. (8) 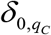 is the Kronecker delta and *n* is the number of elapsed pulses. Terms 1 and 2 are the ^13^C-^1^H and ^15^N-^1^H dipolar couplings, term 3 is the ^13^C and ^15^N CSA, term 4 is the ^1^H chemical shift, and term 5 is the ^15^N-^13^C dipolar coupling.

To apply average Hamiltonian theory (AHT) ^33^, there must be an integer *N* such that 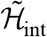 is periodic with period 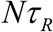. This requires 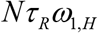,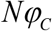, and 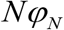 to be integer multiples of 2π. We assume that such an integer *N* exists, that there is no rotary resonance recoupling, i.e. *q_H_ ω*_1,*H*_ + *mω_R_* ≠ 0, and that neither the ^13^C nor ^15^N rf field strengths fulfill Hartman-Hahn matching conditions with the ^1^H rf field, *i.e. q_H_ ω*_1,*H*_τ _R_ + *q_S_φ_S_* ≠ 2*πl*, where *l* is an integer. Under these conditions, the first-order average Hamiltonian is zero because term 4 integrates over 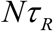 to zero, terms 3 and 5 integrate over each rotor period between pulses to zero, and terms 1 and 2 can also be proven to vanish to first order (**Supporting Information 1**).

For homonuclear polarization transfer by PULSAR, the internal Hamiltonian is identical to the heteronuclear Hamiltonian in Eq. (1) except for an additional “flip-flop” term, 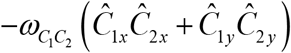. Numerical simulations (**Fig. S3**) indicate that ^13^C-^13^C dipolar couplings have minimal effect on P-TSAR transfer, therefore we neglect the “flip-flop” term in this discussion. A full treatment of the first-order average Hamiltonian of homonuclear dipolar couplings can be found in **Supporting Information 2**.

Because the first-order average Hamiltonian is zero, the second-order average Hamiltonian is expected to dominate. As in CW-TSAR^28, 30^, we attribute P-TSAR transfer to the cross-term between terms 1 and 2 in Eq. (8) in the second-order average Hamiltonian:

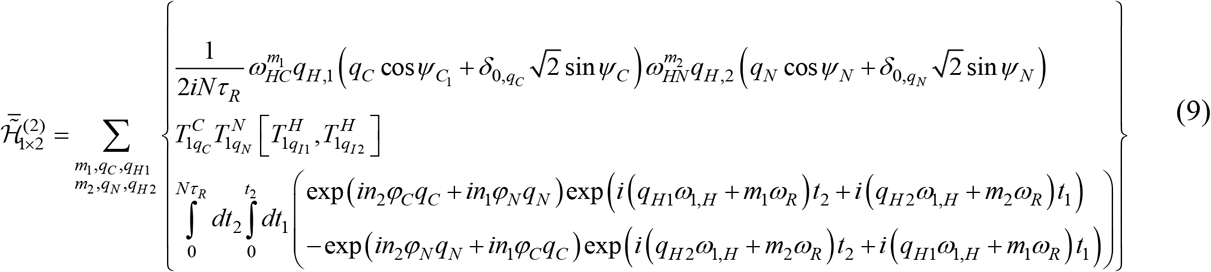

Here *n*_1_ and *n*_2_ are numbers of elapsed pulses in *t_1_* and *t_2_*, and the commutator 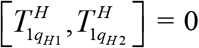 unless *q_H1_* =−*q_H2_*, in which case 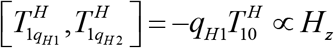. As we show in **Supporting Information 3**, the double integral is zero unless 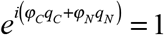 1, or equivalently

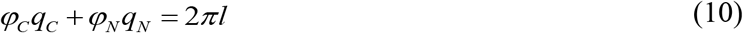

for some integer *l*. Equation (10) can be satisfied in three ways:

1. If the zero-quantum matching condition, *φ_C_* − *φ_N_* = 2*πl* for integer *l*, is met, then when *q_C_* = −*q_N_* = ±1, Eq. (10) will be satisfied. The 1×2 cross term (Eq. 9) produces terms in the second-order average Hamiltonian with spin product operators 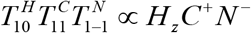 and 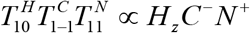. These zero-quantum terms produce polarization transfer from N to C and vice-versa without a change in sign, while the proton acts as a bystander.
2. If the double-quantum matching condition, *φ_C_* + *φ_N_* = 2*π l* for integer *l*, is met, then when *q_C_* = *q_N_* = ±1, Eq. (10) will be satisfied. The 1×2 cross term (Eq. (9)) would produce double-quantum product operators *H_z_C^+^N^+^* and *H_z_C^−^N^−^*. These double-quantum terms produce transfer from N to C and vice-versa, with a change in sign, while the proton acts as a bystander.
3. For any φ_C_ and *φ_N_*, *q_C_* = *q_N_* = 0 satisfies Eq. (10). The spin operator would be 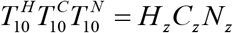. This longitudinal term commutes with both the zero-quantum and double-quantum terms from cases 1 and 2, and will not accomplish any transfer.

A direct analogy can thus be drawn between CW-TSAR and P-TSAR. In both cases, cross terms in the second-order average Hamiltonian between the H–A and H–B dipolar couplings produce double-quantum terms of the form *H_z_A^±^B^±^* or zero-quantum terms of the form *H_z_A^±^B^∓^* in the interaction-frame. In CW-TSAR there are CW spin-locks on all channels, while for P-TSAR it is a pulsed spin-lock for the A and B channels. In both cases, matching conditions on the low-frequency channels dictate when TSAR transfer occurs, and whether it is positive-intensity transfer for the zero-quantum condition or negative-intensity transfer for the double-quantum condition.

### Relation between chemical shift offset, pulse flip angle, and polarization transfer in P-TSAR

The matching conditions in P-TSAR are based on the time-averaged pulsed spin-lock flip angles *φ*_C_ and *φ*_N_. These flip angles are functions of the applied pulse flip angles on each channel and the isotropic offset of each spin. To best design P-TSAR experiments, we now consider how the choices of pulse flip angle and rf carrier frequencies dictate the chemical shifts that survive the pulsed spin-lock and fulfill the zero-quantum or double-quantum matching conditions.

Consider a single spin at a chemical shift offset Ω subjected to a train of rf pulses with flip angle *θ* about the *x*-axis at the center of each rotor period. This spin undergoes three successive rotations per rotor period (**Fig. 6a**): a rotation about the *z*-axis during the two free evolution periods (1 to 2 and 3 to 4), sandwiching a *θ* rotation about the *x*-axis during the pulse (2 to 3). These three successive rotations can be expressed as:

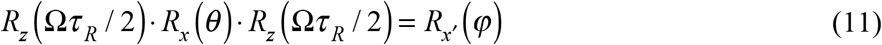

**Figure 6.**
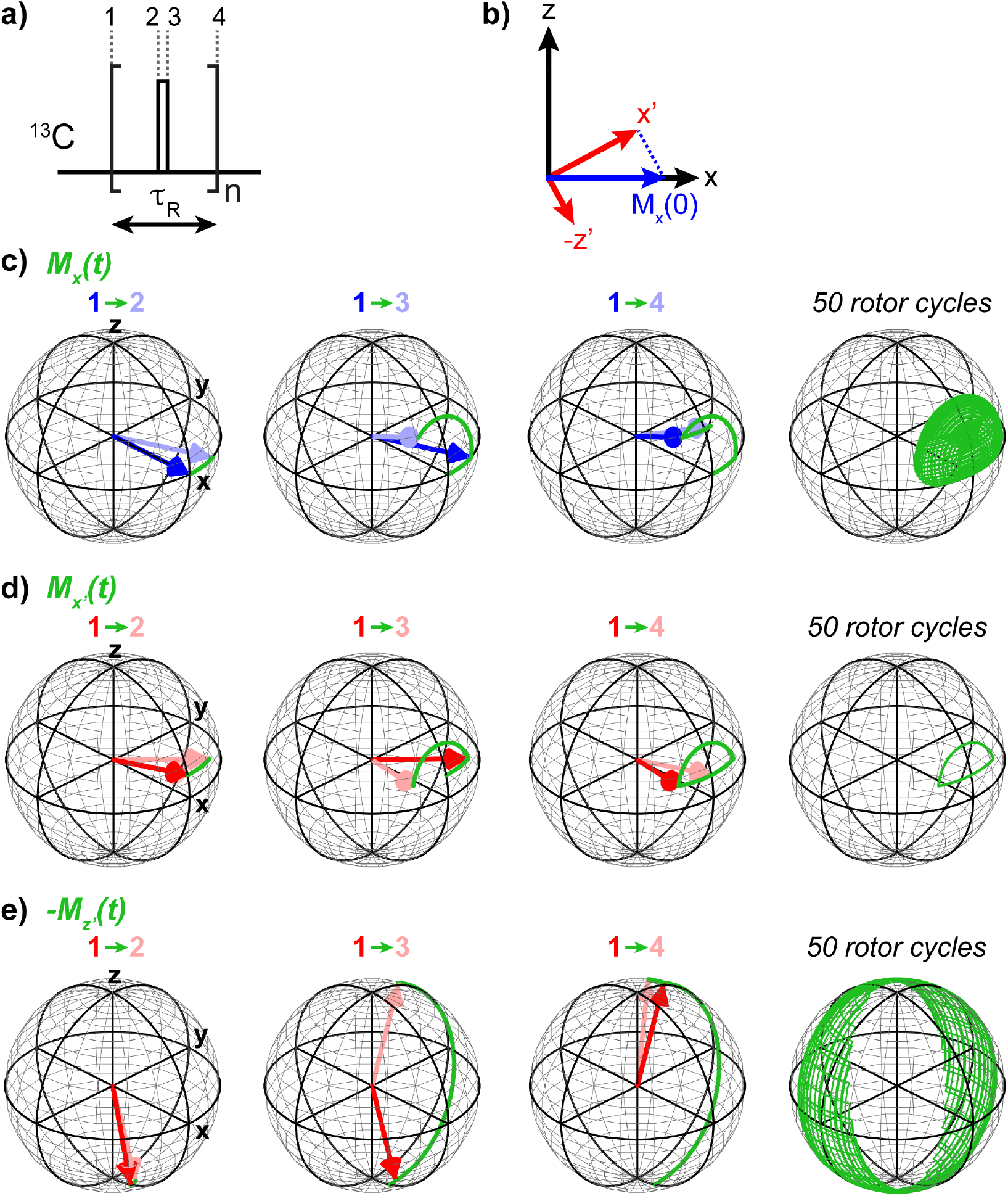
Magnetization trajectories under pulsed spin-lock. **(a)** Diagram of the basic pulsed spin-lock unit. The free evolution periods of 1→2 and 3→4 rotate the magnetization around the *z*-axis due to the isotropic chemical shift offset, while the rf pulse during 2→3 rotates the magnetization around the *x*-axis. The pulsed spin-lock axis is tilted from the *x*-axis in the *xz*-plane. The initial *x*-magnetization, *M_x_*(0), has two components along *x’* and *z’*. (**c-e**). Magnetization trajectories under pulsed spin-lock, calculated in MATLAB using an rf field of 50 kHz, a pulse flip angle of 90°, and a chemical shift offset of 17.5 ppm Trajectory of *x*-magnetization. The initial and final locations are not at the same spot on the Bloch sphere. **(d)** Trajectory of *M_x_*’. This magnetization begins and ends the rotor period at the same position on the Bloch sphere, hence it is spin-locked. **(e)** Trajectory of *M_z_*’. This magnetization traces out a broad trajectory, indicating rapid decay.

The net effect of the three rotations is equivalent to a single rotation of angle *φ* about the pulsed spin-lock axis 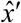 (**Fig. 6b**), which can be defined as the rotation operator 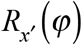. This rotation is repeated many times during the pulsed spin-lock period (**Fig. 6c**), so that only magnetization along the spin-lock axis will be fully retained **(Fig. 6d**) while magnetization that is perpendicular to the effective spin-lock axis will precess to a different position on the Bloch sphere at the end of each rotor period (**Fig. 6e**). Any magnetization that is not parallel to the 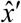 axis thus decays rapidly and can be discarded. Placing the spin-lock pulses in the center of each rotor period ensures that the spin-lock axis lies in the *xz*-plane; any initial *y*-magnetization is dephased during the spin-lock, thus allowing quadrature detection of the indirect dimension. However, *x’* is not necessarily along *x*, but is instead in the *xz*-plane, therefore only a portion of the initial *x-*magnetization is properly spin-locked while the rest is dephased:

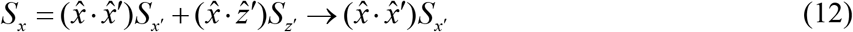

The initial *x*-magnetization, M_*x*_, can be expressed in the *x’* and *z’* basis by projection (**Fig. 6b**). The time-dependent trajectory of M_*x*_ can then be expressed as two oscillating *x’* and *z’* components:

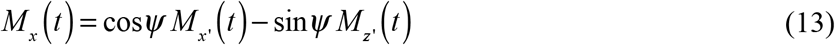

Magnetization along *z’* is perpendicular to the spin-lock axis and decays rapidly. A pulsed spin-lock will scale down magnetization by a factor of 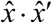, with a different direction of 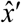 for each isochromat (**Fig. S4**). This “read-in” loss is followed by a “read-out” loss in which magnetization is projected from the spin-lock direction back to the *x*-axis for detection (**Fig. 7a**). Treating the rf pulse as finite rather than a δ-function pulse gives a more accurate pulsed spin-lock profile (**Fig. 7b, Supporting Information 4**). For small pulse flip angles, the “read-in” and “read-out” losses are minimal for spins on resonance or at an rf offset near *nω_r_*. For pulse flip angles near 360°, losses are minimal on resonance and at an rf offset of about 1.5*ω_r_*. The pulsed spin-lock bandwidth increases as the nominal pulse flip angle increases from 0° to 180° or decreases from 360° to 180°.

**Figure 7.**
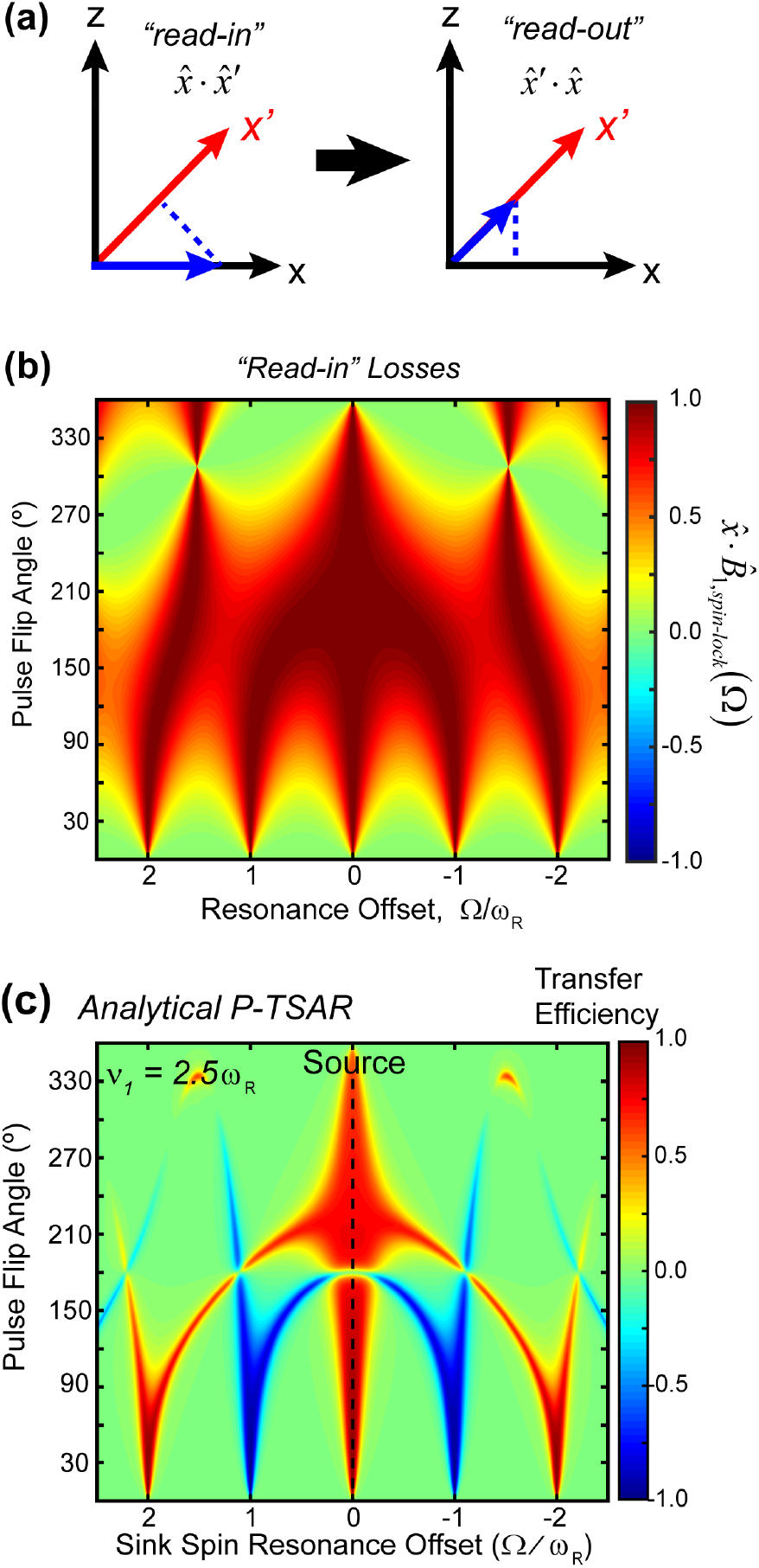
“Read-in” and “read-out” losses associated with a pulsed spin-lock. **(a)**“Reading in” and “reading out” of the pulsed spin-lock causes magnetization losses due to projection of one axis onto the other. **(b)** Contour plot showing the “read-in” losses as a function of pulse flip angle and resonance offset. 50 kHz rf pulses were used in these calculations. **(c)** Calculated P-TSAR efficiencies as a function of the sink-spin offset. The source spin (C1) was placed on resonance. Positive transfer occurs at the zero-quantum condition (φC1 = φC2) while negative transfer occurs at the double-quantum condition (φC1 + φC2 = 2π).

This analytical model not only gives the pulsed spin-lock axis, but also the net rotation, *φ*, during each rotor period. For visualization purposes, we can convert this *φ* into a time-averaged pulsed spin-lock field strength *ω_SL_*:

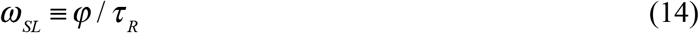

This time-averaged spin-lock field is analogous to the CW spin-lock field in PAR and ^PAIN^CP. From the average Hamiltonian analysis, the P-TSAR sequences have double- and zero-quantum matching conditions 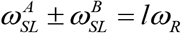 for integer values of *l*. The zero-quantum condition leads to positive transfer intensities while the double-quantum condition leads to negative transfer intensities. This differs from typical implementations of CW-TSAR experiments, which only fulfill zero-quantum matching conditions.

To model the P-TSAR transfer, we combine the “read-in” and “read-out” losses with the matching conditions: the overall transfer efficiency *η*_*A*_→_*B*_ from spin *A* to *B* is the product of three factors: the “read-in” loss 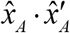, projection of one spin’s spin-lock axis onto the other, 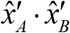, and the “read-out” loss 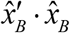. These three factors are further multiplied by Lorentzian functions centered at the zero- and double-quantum matching conditions. Finally, we include a scaling factor (1− *τ_p_*/τ_R_) to account for the finite pulse length, which reduces the effective transfer time for long pulses:

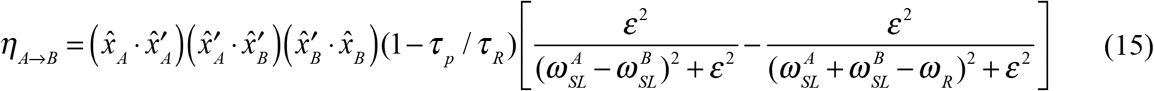

Eq. (15) was used to calculate the polarization transfer efficiencies (**Fig. 7c, 8c**) with a Lorentzian width ε=0.05ω_r_. This width was chosen empirically by varying *ε* in the analytical model until the local bandwidth is consistent with numerical simulations and experiments. The resulting polarization transfer profile (**Fig. 8c**) has a nearly identical chemical shift and flip-angle dependence as that obtained from numerical simulation (**Fig. 8b**). Positive transfer is maximized when the source and sink spins have the same absolute offset. Negative transfer is obtained when the sink spin is ~ *ω_r_* away from the carrier for small flip angles, but splits into two separate branches for larger flip angles. A nominal flip angle of 180° gives vanishing transfer. With flip angles larger than 180°, the bandwidth for positive zero-quantum transfer is significantly larger than for smaller flip angles.

**Figure 8.**
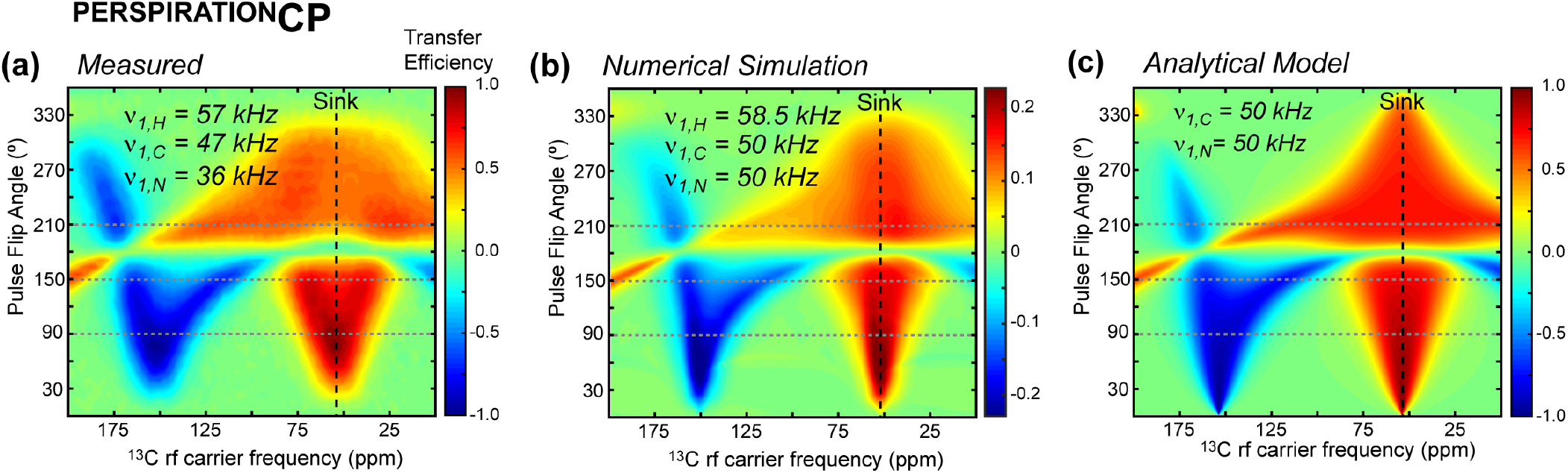
Measured, simulated, and calculated pulsed spin-lock polarization transfer efficiencies as a function of the ^13^C rf carrier frequency and the pulse flip angle. The experiments were conducted on an 800 MHz spectrometer under 20 kHz MAS. The calculations were conducted for the same conditions. **(a)** Measured ^PERSPIRATION^CP efficiencies of f-MLF as a function of ^13^C carrier frequency and ^13^C pulse flip angle. The 2D heat map is generated from 25 x 80 1D spectra measured as a function of ^13^C pulsed-spin-lock flip angles (0° to 360° in 15° increments) and rf carrier frequencies (0 to 200 ppm in 2.5 ppm increments.) The integrated Cθ band intensities (46 ppm to 59 ppm) of the spectra are represented in the plot. The rf field strengths of the ^PERSPIRATION^CP block are 57 kHz, 47 kHz, and 36 kHz for ^1^H, ^13^C and ^15^N, respectively. **(b)** Numerical simulations of ^PERSPIRATION^CP polarization transfer from an on-resonance ^15^N to ^13^Cα. **(c)**^PERSPIRATION^CP transfer efficiencies calculated using the analytical model.

To test the isotropic shift dependence of P-TSAR, we carried out a pseudo-3D ^PERSPIRATION^CP experiment on the model peptide formyl-Met-Leu-Phe (f-MLF), in which we varied the ^13^C rf carrier frequency and pulse flip angle in the two indirect dimensions. **Fig. 8a** plots the integrated intensity of the three Cα peaks as a function of these two parameters (**Fig. S5**). Remarkably, the experiment gives the same qualitative features as predicted in the numerical simulations and the analytical model. Maximum positive transfer occurs when the source and sink spin chemical shift offsets are both small, while maximum negative transfer occurs when the ^13^C carrier is about *ω*_*r*_ away from the sink spins for small pulse flip angles.

## Discussion

Our first implementation of the heteronuclear ^PERSPIRATION^CP experiment (**Fig. S1d**) ^34^ placed the spin-lock pulses at the end of each rotor period. This timing refocuses magnetization of spins that are on resonance or at ~*nω_r_* from the carrier frequency for small flip angles, but does not refocus other off-resonance magnetization. Off-resonance signals have both *x-* and *y*-components at the end of the pulsed spin-lock period, which require a *z*-filter before and after this period to obtain clean, in-phase *x*-magnetization. Omitting these *z*-filters results in spectra with severe phase distortion. Inclusion of the *z*-filters leads to significant “read-in” and “read-out” losses by discarding the *y*-component (**Fig. S5a**). Spins that are ~0.5*ω_r_* off-resonance for small flip angles do not survive the pulsed spin-lock and therefore do not contribute to polarization transfer (**Fig. S5b,c**). ^PERSPIRATION^CP with pulses at the end of each rotor period results in low transfer efficiency above 60 ppm and below 20 ppm and no transfer for aromatic signals at 110-140 ppm. Centering the spin-lock pulses in the rotor period refocuses magnetization along an axis in the *xz*-plane. This allows us to remove the *z*-filters, which facilitates quadrature detection and broadens the bandwidth of these experiments.

Previous studies have shown that a pulsed spin-lock can extend the coherence lifetime in solids by about 10-fold compared to a spin echo ^32, 46^.These past implementations of pulsed spin-locks to solids used simple spin systems, such as CaF_2_ ^47^ in the absence of MAS, or amino acids with a single ^13^C label ^46^, in which spectral resolution and bandwidth were not of primary concern. In this study, we demonstrate the use of a pulsed spin-lock in the presence of significant isotropic chemical shift dispersion, wherein maximizing the pulsed spin-lock bandwidth is of utmost importance to enable the identification of as many long-range contacts as possible. By placing a pulse in the center of each rotor-period, we make the sideband spacing significantly larger compared to previous MAS pulsed spin-lock experiments that pulsed only every 20 rotor periods for similar spinning frequencies ^46^. Moreover, our careful choice of the pulse flip angle maximizes the breadth of the envelope of magnetization that survives the pulsed spin-lock compared to experiments that use flip angles less than 90° in order to maximize the relaxation time constant. Importantly, in contrast to previous work that uses a pulsed spin-lock to extend coherence time for greater sensitivity^32, 46–49^, here we use a pulsed spin-lock to generate distinct average Hamiltonians along the pulsed spin-lock directions that accomplish internuclear transfer.

The excellent agreement between our analytical model (**Fig. 8c**), numerical simulations (**Fig. 8b**), and measured (**Fig. 8a**) polarization transfer profiles indicates that once a ^1^H CW field strength is chosen, the main factors dictating polarization transfer are the pulse flip angle and the chemical shift offset that fulfill the zero- or double-quantum matching conditions, Eq. (10). These matching conditions are well-predicted by our analytical model, hence the ^13^C flip angle and carrier frequency can be chosen by inspecting the calculated analytical 2D map for a given magnetic field strength and MAS frequency (**Appendix S3**). Once the ^13^C carrier frequency and pulse flip angle are chosen, PULSAR can be optimized with only two steps. First, we optimize the ^1^H CW rf field with a 1D experiment that includes a Cα selective echo before the P-TSAR block, by maximizing the non-Cα intensities (**Fig. S1a**). Second, we locally optimize the ^13^C pulse length to again maximize the transferred intensity in the desired spectral regions. ^PERSPIRATION^CP experiments require an additional step of optimizing the ^15^N pulse length with respect to the ^13^C pulse length, which is simple once the ^13^C flip angle has been selected. Importantly, these optimization steps are uncoupled and can be run independent of each other.

This optimization routine is much faster and easier than the multi-parameter optimization for CW-TSAR experiments, as a result of the extreme sensitivity of PAR and ^PAIN^CP’s matching conditions to rf field strengths, spin geometry, and ^1^H density. Even for a given sample, different spin pairs have different optimal rf field strengths. Moreover, the rf field strengths that result in good polarization transfer for CW-TSAR experiments are inter-dependent. Such multidimensional parameter searches are very time-consuming, especially for 2D correlation experiments. For example, a recent investigation involving the 2D ^15^N-^15^N PAR experiment conducted 29 × 29 = 841 separate 2D’s to search for the optimal ^1^H and ^15^N CW spin-lock field strengths ^50^. Simulations can facilitate this parameter search, but often show significant discrepancies from experiments ^31^. The optimized parameters can also change significantly between samples, which make it difficult to implement CW-TSAR experiments on low-sensitivity biological samples. In comparison, P-TSAR experiments do not require sample-specific optimization; the matching conditionals are easily predicted using analytical theory, and the user can choose the pulse flip angle and rf carrier frequency in advance. Moreover, the ^1^H CW field strength in P-TSAR experiments is only weakly coupled to other parameters, which further simplifies the optimization of P-TSAR experiments.

## Conclusions

We have introduced two pulsed third-spin-assisted recoupling (P-TSAR) experiments that efficiently transfer polarization between homonuclear and heteronuclear spin pairs using a low rf duty cycle. The PULSAR and the improved ^PERSPIRATION^CP experiments are demonstrated on GB1 and glucagon amyloid fibrils and give rise to many long-range correlations that constrain the 3D structures of proteins. Using numerical simulations and analytical theory, we found that the first-order average Hamiltonian for the P-TSAR experiments is averaged to zero, apart from a homonuclear dipolar coupling term that has negligible effects on P-TSAR. We found that the second-order average Hamiltonian yields non-zero cross terms between the ^1^H-^13^C_1_ (^1^H-^15^N) and ^1^H-^13^C_2_ (^1^H-^13^C) dipolar couplings, which drive polarization transfer. These properties are the same as the PAR and ^PAIN^CP experiments. P- and CW-TSAR experiments give similar long-range contacts, but P-TSAR experiments use 5-10 fold less rf energy than their CW counterparts, and are much easier to optimize. The current study demonstrates these P-TSAR experiments on 600 and 800 MHz spectrometers under 20 kHz MAS, but the methodology can be extended to other magnetic field strengths, MAS frequency regimes, and spin pairs.

## Acknowledgements

This work is supported by NIH grant AG059661 to M.H. M.D.G. is supported by an NIH Ruth L. Kirschstein Individual National Research Service Award (1F31AI133989).

---

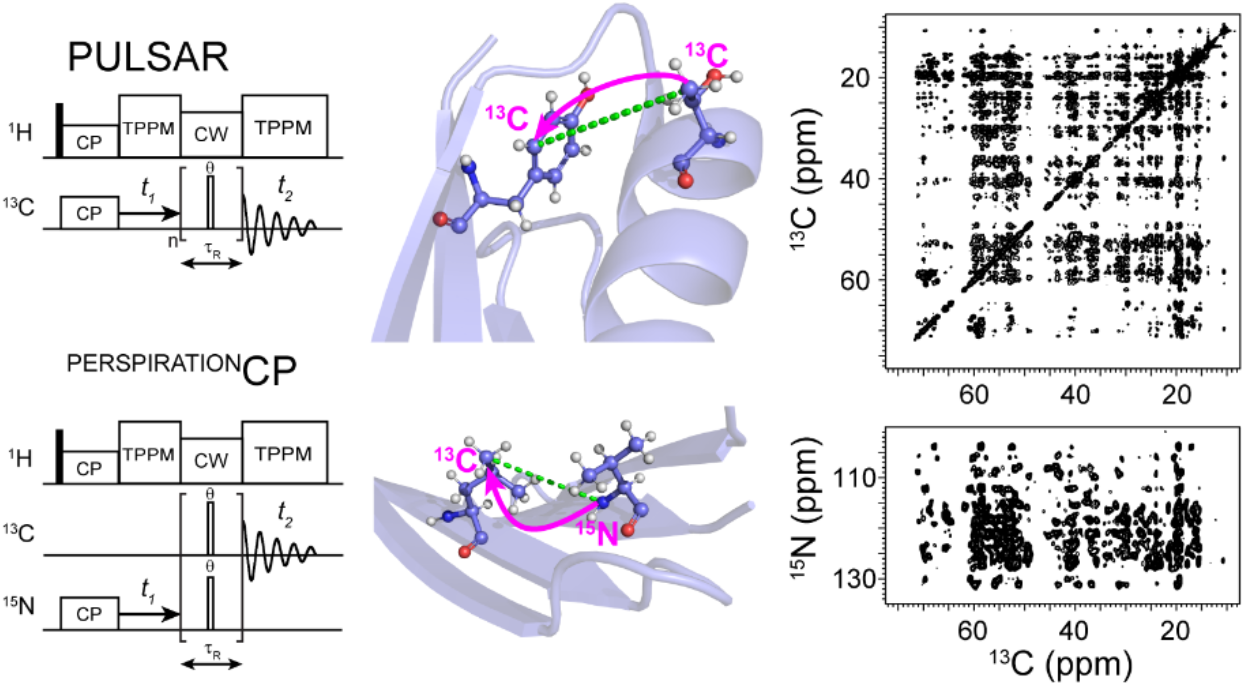

## References

1. Wang, S. L.; Munro, R. A.; Shi, L. C.; Kawamura, I.; Okitsu, T.; Wada, A.; Kim, S. Y.; Jung, K. H.; Brown, L. S.; Ladizhansky, V., Solid-state NMR spectroscopy structure determination of a lipid-embedded heptahelical membrane protein. Nat Methods 2013, 10 (10), 1007–1012.

2. Möbius, K.; Kazemi, S.; Güntert, P.; Jakob, A.; Heckel, A.; Becker-Baldus, J.; Glaubitz, C., Global response of diacylglycerol kinase towards substrate binding observed by 2D and 3D MAS NMR. Sci. Rep. 2019, 9 (1), 3995.

3. Jekhmane, S.; Medeiros-Silva, J.; Li, J.; Kümmerer, F.; Müller-Hermes, C.; Baldus, M.; Roux, B.; Weingarth, M., Shifts in the selectivity filter dynamics cause modal gating in K+ channels. Nat. Commun. 2019, 10 (1), 123.

4. Mandala, V. S.; Loftis, A. R.; Shcherbakov, A. A.; Pentelute, B. L.; Hong, M., Atomic Structures of Closed and Open Influenza B M2 Proton Channel Reveal the Conduction Mechanism. Nat. Struc. Mol. Biol. 2020, 27, 160–167.

5. Paravastu, A. K.; Leapman, R. D.; Yau, W.-M.; Tycko, R., Molecular structural basis for polymorphism in Alzheimer’s β-amyloid fibrils. Proc. Natl. Acad. Sci. USA 2008, 105 (47), 18349–18354.

6. Wasmer, C.; Lange, A.; Van Melckebeke, H.; Siemer, A. B.; Riek, R.; Meier, B. H., Amyloid fibrils of the HET-s(218-289) prion form a beta solenoid with a triangular hydrophobic core. Science 2008, 319, 1523–1526.

7. Bertini, I.; Gonnelli, L.; Luchinat, C.; Mao, J.; Nesi, A., A New Structural Model of Aβ40 Fibrils. J. Am. Chem. Soc. 2011, 133 (40), 16013–16022.

8. Lu, J. X.; Qiang, W.; Yau, W. M.; Schwieters, C. D.; Meredith, S. C.; Tycko, R., Molecular structure of β-amyloid fibrils in Alzheimer’s disease brain tissue. Cell 2013, 154, 1257–68.

9. Colvin, M. T.; Silvers, R.; Ni, Q. Z.; Can, T. V.; Sergeyev, I.; Rosay, M.; Donovan, K. J.; Michael, B.; Wall, J.; Linse, S.; Griffin, R. G., Atomic Resolution Structure of Monomorphic Aβ42 Amyloid Fibrils. J. Am. Chem. Soc. 2016, 138 (30), 9663–9674.

10. Wälti, M. A.; Ravotti, F.; Arai, H.; Glabe, C. G.; Wall, J. S.; Böckmann, A.; Güntert, P.; Meier, B. H.; Riek, R., Atomic-resolution structure of a disease-relevant Aβ(1–42) amyloid fibril. Proc. Natl. Acad. Sci. USA 2016, 113 (34), E4976–E4984.

11. Tuttle, M. D.; Comellas, G.; Nieuwkoop, A. J.; Covell, D. J.; Berthold, D. A.; Kloepper, K. D.; Courtney, J. M.; Kim, J. K.; Barclay, A. M.; Kendall, A.; Wan, W.; Stubbs, G.; Schwieters, C. D.; Lee, V. M. Y.; George, J. M.; Rienstra, C. M., Solid-state NMR structure of a pathogenic fibril of full-length human α-synuclein. Nat. Struct. Mol. Biol. 2016, 23, 409.

12. Gelenter, M. D.; Smith, K. J.; Liao, S. Y.; Mandala, V. S.; Dregni, A. J.; Lamm, M. S.; Tian, Y.; Xu, W.; Pochan, D. J.; Tucker, T. J.; Su, Y.; Hong, M., The peptide hormone glucagon forms amyloid fibrils with two coexisting b-strand conformations. Nat. Struc. Mol. Biol 2019, 26, 592–598.

13. Dregni, A. J.; Mandala, V. S.; Wu, H.; Elkins, M. R.; Wang, H. K.; Hung, I.; DeGrado, W. F.; Hong, M., In vitro 0N4R tau fibrils contain a monomorphic b-sheet core enclosed by dynamically heterogeneous fuzzy coat segments. Proc. Natl. Acad. Sci. U.S.A 2019, 116, 16357–16366.

14. Phyo, P.; Gu, Y.; Hong, M., Impact of acidic pH on plant cell wall polysaccharide structure and dynamics: insights into the mechanism of acid growth in plants from solid-state NMR. Cellulose 2019, 26 (1), 291–304.

15. Wang, T.; Park, Y. B.; Caporini, M. A.; Rosay, M.; Zhong, L.; Cosgrove, D. J.; Hong, M., Sensitivity-enhanced solid-state NMR detection of expansin’s target in plant cell walls. Proc. Natl. Acad. Sci. U. S. A. 2013, 110, 16444–16449.

16. Kang, X.; Kirui, A.; Muszyński, A.; Widanage, M. C. D.; Chen, A.; Azadi, P.; Wang, P.; Mentink-Vigier, F.; Wang, T., Molecular architecture of fungal cell walls revealed by solid-state NMR. Nat. Commun. 2018, 9 (1), 2747.

17. Romaniuk, J. A. H.; Cegelski, L., Peptidoglycan and Teichoic Acid Levels and Alterations in Staphylococcus aureus by Cell-Wall and Whole-Cell Nuclear Magnetic Resonance. Biochemistry 2018, 57 (26), 3966–3975.

18. Bennett, A. E.; Rienstra, C. M.; Griffiths, J. M.; Zhen, W.; Jr., P. T. L.; Griffin, R. G., Homonuclear radio frequency-driven recoupling in rotating solids. J. Chem. Phys. 1998, 108 (22), 9463–9479.

19. Gullion, T.; Schaefer, J., Rotational-echo double-resonance NMR. J. Magn. Reson. 1989, 81 (1), 196–200.

20. Hing, A. W.; Vega, S.; Schaefer, J., Transferred-echo double-resonance NMR. J. Magn. Reson. 1992, 96 (1), 205–209.

21. Jaroniec, C. P.; Filip, C.; Griffin, R. G., 3D TEDOR NMR Experiments for the Simultaneous Measurement of Multiple Carbon−Nitrogen Distances in Uniformly ^13^C,^15^N-Labeled Solids. J. Am. Chem. Soc. 2002, 124 (36), 10728–10742.

22. Bloembergen, N., On the interaction of nuclear spins in a crystalline lattice. Physica 1949, 15 (3), 386–426.

23. Takegoshi, K.; Nakamura, S.; Terao, T., ^13^C–^1^H dipolar-assisted rotational resonance in magic-angle spinning NMR. Chem. Phys. Lett. 2001, 344 (5), 631–637.

24. Morcombe, C. R.; Gaponenko, V.; Byrd, R. A.; Zilm, K. W., Diluting abundant spins by isotope edited radio frequency field assisted diffusion. J. Am. Chem. Soc. 2004, 126, 7196–7197.

25. Hou, G.; Yan, S.; Trébosc, J.; Amoureux, J.-P.; Polenova, T., Broadband homonuclear correlation spectroscopy driven by combined R2_n_^v^ sequences under fast magic angle spinning for NMR structural analysis of organic and biological solids. J. Magn. Reson. 2013, 232, 18–30.

26. Demers, J.-P.; Chevelkov, V.; Lange, A., Progress in correlation spectroscopy at ultra-fast magic-angle spinning: Basic building blocks and complex experiments for the study of protein structure and dynamics. Solid State Nucl. Magn. Reson. 2011, 40 (3), 101–113.

27. Paul, S.; Takahashi, H.; Hediger, S.; De Paëpe, G., Chapter Three - Third Spin-Assisted Recoupling in SSNMR: Theoretical Insights and Practicable Application to Biomolecular Structure Determination. In Annu. Rep. NMR Spectrosc., Webb, G. A., Ed. Academic Press: 2015; Vol. 85, pp 93–142.

28. De Paëpe, G.; Lewandowski, J. R.; Loquet, A.; Böckmann, A.; Griffin, R. G., Proton assisted recoupling and protein structure determination. J. Chem. Phys. 2008, 129 (24), 245101.

29. Lewandowski, J. R.; De Paëpe, G.; Griffin, R. G., Proton Assisted Insensitive Nuclei Cross Polarization. J. Am. Chem. Soc. 2007, 129 (4), 728–729.

30. De Paëpe, G.; Lewandowski, J. R.; Loquet, A.; Eddy, M.; Megy, S.; Böckmann, A.; Griffin, R. G., Heteronuclear proton assisted recoupling. J. Chem. Phys. 2011, 134 (9), 095101.

31. Donovan, K. J.; Jain, S. K.; Silvers, R.; Linse, S.; Griffin, R. G., Proton-Assisted Recoupling (PAR) in Peptides and Proteins. J. Phys. Chem. B 2017, 121 (48), 10804–10817.

32. Ostroff, E. D.; Waugh, J. S., Multiple Spin Echoes and Spin Locking in Solids. Phys. Rev. Lett. 1966, 16 (24), 1097–1098.

33. Haeberlen, U.; Waugh, J. S., Coherent Averaging Effects in Magnetic Resonance. Phys. Rev. 1968, 175 (2), 453–467.

34. Gelenter, M. D.; Hong, M., Efficient 15N-13C Polarization Transfer by Third-Spin-Assisted Pulsed Cross-Polarization Magic-Angle-Spinning NMR for Protein Structure Determination. J. Phys. Chem. B 2018, 122, 8367–8379.

35. Franks, W. T.; Zhou, D. H.; Wylie, B. J.; Money, B. G.; Graesser, D. T.; Frericks, H. L.; Sahota, G.; Rienstra, C. M., Magic-angle spinning solid-state NMR spectroscopy of the beta1 immunoglobulin binding domain of protein G (GB1): 15N and 13C chemical shift assignments and conformational analysis. J. Am. Chem. Soc. 2005, 127, 12291–12305.

36. Gelenter, M. D.; Wang, T.; Liao, S.-Y.; O’Neill, H.; Hong, M., ^2^H–^13^C correlation solid-state NMR for investigating dynamics and water accessibilities of proteins and carbohydrates. J. Biomol. NMR 2017, 68 (4), 257–270.

37. Mandala, V. S.; Liao, S.-Y.; Gelenter, M. D.; Hong, M., The Transmembrane Conformation of the Influenza B Virus M2 Protein in Lipid Bilayers. Sci. Rep. 2019, 9 (1), 3725.

38. Bernard, G. M.; Goyal, A.; Miskolzie, M.; McKay, R.; Wu, Q.; Wasylishen, R. E.; Michaelis, V. K., Methylammonium lead chloride: A sensitive sample for an accurate NMR thermometer. J. Magn. Reson. 2017, 283, 14–21.

39. Bennett, A. E.; Rienstra, C. M.; Auger, M.; Lakshmi, K. V.; Griffin, R. G., Heteronuclear decoupling in rotating solids. J. Chem. Phys. 1995, 103 (16), 6951–6958.

40. Lee, W.; Tonelli, M.; Markley, J. L., NMRFAM-SPARKY: enhanced software for biomolecular NMR spectroscopy. Bioinformatics 2015, 31 (8), 1325–1327.

41. Güntert, P.; Mumenthaler, C.; Wüthrich, K., Torsion angle dynamics for NMR structure calculation with the new program Dyana. J. Mol. Biol. 1997, 273 (1), 283–298.

42. Shen, Y.; Bax, A., Protein backbone and sidechain torsion angles predicted from NMR chemical shifts using artificial neural networks. J. Biomol. NMR 2013, 56 (3), 227–241.

43. Veshtort, M.; Griffin, R. G., SPINEVOLUTION: A powerful tool for the simulation of solid and liquid state NMR experiments. J. Magn. Reson. 2006, 178 (2), 248–282.

44. Wylie, B. J.; Sperling, L. J.; Nieuwkoop, A. J.; Franks, W. T.; Oldfield, E.; Rienstra, C. M., Ultrahigh resolution protein structures using NMR chemical shift tensors. Proc. Natl. Acad. Sci. USA 2011, 108 (41), 16974–16979.

45. Gratzer, W. B.; Bailey, E.; Beaven, G. H., Conformational states of glucagon. Biochem. Biophys. Res. Commun. 1967, 28, 914–919.

46. Petkova, A. T.; Tycko, R., Sensitivity Enhancement in Structural Measurements by Solid State NMR through Pulsed Spin Locking. J. Magn. Reson. 2002, 155 (2), 293–299.

47. Rhim, W. K.; Burum, D. P.; Elleman, D. D., Multiple-Pulse Spin Locking in Dipolar Solids. Phys. Rev. Lett. 1976, 37 (26), 1764–1766.

48. Suwelack, D.; Waugh, J. S., Quasistationary magnetization in pulsed spin-locking experiments in dipolar solids. Phys. Rev. B 1980, 22 (11), 5110–5114.

49. Rhim, W. K.; Burum, D. P.; Elleman, D. D., Calculation of spin–lattice relaxation during pulsed spin locking in solids. J. Chem. Phys. 1978, 68 (2), 692–695.

50. Donovan, K. J.; Silvers, R.; Linse, S.; Griffin, R. G., 3D MAS NMR Experiment Utilizing Through-Space ^15^N–^15^N Correlations. J. Am. Chem. Soc. 2017, 139 (19), 6518–6521.

