## Supporting Information for "Pulsed Third-Spin-Assisted Recoupling NMR for Obtaining Long-Range ^13^C-^13^C and ^15^N-^13^C Distance Restraints"

**1. Analysis of the H-X heteronuclear dipolar coupling terms in the first-order average Hamiltonian of P-TSAR experiments**

Below we explicitly show that the first-order average Hamiltonian for the H-X heteronuclear dipolar couplings (terms 1 and 2) in Eq. (**8**) are zero. The first-order average Hamiltonian is calculated as:

$$\bar{\tilde{\mathcal{H}}}^{(1)} = \frac{1}{N\tau_R} \int_0^{N\tau_R} \tilde{\mathcal{H}}(t) dt \tag{S1}$$

Integrating the first-order average Hamiltonian for term 1 in Eq. **(8)**, or equivalently term 2 if all '*C*' are replaced with '*N*', we obtain:

$$\bar{\tilde{\mathcal{H}}}_1^{(1)} = \sum_{\substack{m_1=-2 \\ m_1\neq 0}}^{2} \sum_{q_{C=-1}}^{1} \sum_{\substack{q_{H1}=-1 \\ q_{H1}\neq 0}}^{1} \frac{\omega_{HC}^{m} q_{H1}\left[q_C \cos(\psi_C) + \delta_{0,q_C}\sqrt{2}\sin(\psi_C)\right] T_{1q_{H1}}^{H} T_{1q_C}^{C}}{N\tau_R} \int_0^{N\tau_R} \exp\left(in\varphi_C q_C\right) \exp\left(i[q_{H1}\omega_{1,H} + m\omega_R]t\right) dt \tag{S2}$$

where $n \equiv \lfloor t/\tau_R + 1/2 \rfloor$. Disregarding terms outside the integral, we first separate the integral over *N* rotor periods into a sum of integrals of one rotor period each, thereby removing the *t* dependence of *n*:

$$\begin{aligned} \bar{\tilde{\mathcal{H}}}_1^{(1)} &\propto \int_0^{N\tau_R} \exp\left(in\varphi_C q_C\right) \exp\left(i[q_{H1}\omega_{1,H} + m\omega_R]t\right) dt \\ &= \sum_{n=0}^{N-1} \exp\left(in\varphi_C q_C\right) \int_{(n-\frac{1}{2})\tau_R}^{(n+\frac{1}{2})\tau_R} \exp\left(i[q_{H1}\omega_{1,H} + m\omega_R]t\right) dt \end{aligned} \tag{S3}$$

We have combined the first integral, $t \in [0, \frac{\tau_R}{2}]$ with the final one, $t \in [(N-\frac{1}{2})\tau_R, N\tau_R]$, by noting that all time dependent terms are $N\tau_R$ periodic. We assume that we avoid fulfilling the rotary resonance condition, $q_H\omega_{1,H} + m\omega_R \neq 0$, and as such can explicitly integrate:

$$\begin{aligned} \bar{\tilde{\mathcal{H}}}_1^{(1)} &\propto \sum_{n=0}^{N-1} \exp\left(in\varphi_C q_C + inq_{H1}\omega_{1,H}\tau_R\right)(-1)^m \frac{\exp(+iq_{H1}\omega_{1,H}\frac{\tau_R}{2}) - \exp(-iq_{H1}\omega_{1,H}\frac{\tau_R}{2})}{i[q_{H1}\omega_{1,H} + m\omega_R]} \\ &= \frac{2(-1)^m \sin(q_H\omega_{1,H}\frac{\tau_R}{2})}{q_{H1}\omega_{1,H} + m\omega_R} \sum_{n=0}^{N-1} \left(e^{i\left[\varphi_C q_C + q_H\omega_{1,H}\tau_R\right]}\right)^n \end{aligned} \tag{S4}$$

The summation is zero unless $e^{i\left[\varphi_C q_C + q_H\omega_{1,H}\tau_R\right]} = 1$, as it is a sum over roots of unity. This can be seen by noting that a complex exponential is a point on the unit circle, and these *N* complex exponentials are equally spaced about the unit circle and end up at 1: either they are all 1 or they cancel exactly by symmetry. The case $e^{i\left[\varphi_C q_C + q_H\omega_{1,H}\tau_R\right]} = 1$ is effectively the Hartman-Hahn matching condition, which we have assumed is not fulfilled, to be able to apply the Average Hamiltonian Theory (AHT). Therefore, under the conditions of the P-TSAR experiments, the first-order average Hamiltonian for terms 1 and 2 is zero.

**2. Interaction-frame transformation and the first-order average Hamiltonian of the homonuclear zero-quantum flip-flop term**

We wish to transform the flip-flop portion of the homonuclear dipolar coupling, $\mathcal{H}_{CCZQ} = -\omega_{C_1C_2}\left(\hat{C}_{1x}\hat{C}_{2x} + \hat{C}_{1y}\hat{C}_{2y}\right) = \omega_{C_1C_2}(T_{11}^{C_1}T_{1-1}^{C_2} + T_{1-1}^{C_1}T_{11}^{C_2})$ into the interaction frame of the rf pulses and isotropic shifts for both $C_1$ and $C_2$. Following the examples for $\hat{C}_z$ in the main text, Eq. (**5-7**), we first rotate $\mathcal{H}_{CCZQ}$ by $-\Omega_{C_1}(t-n\tau_R)$ and $-\Omega_{C_2}(t-n\tau_R)$ about $\hat{C}_{1z}$ and $\hat{C}_{2z}$, respectively; then rotate it by $-\psi_{C_1}$ and $-\psi_{C_2}$ about $\hat{C}_{1y}$ and $\hat{C}_{2y}$, respectively. These rotations convert $\hat{C}_{1x}$ to $\hat{C}_{1x'}$ and $\hat{C}_{2x}$ to $\hat{C}_{2x'}$. We then rotate $\mathcal{H}_{CCZQ}$ again by -90° about $\hat{C}_{1y}$ and $\hat{C}_{2y}$ to make the spin-lock axes $\hat{C}_{1z'}$ and $\hat{C}_{2z'}$ instead of $\hat{C}_{1x'}$ and $\hat{C}_{2x'}$. We finally rotate $\mathcal{H}_{CCZQ}$ by $-n\varphi_{C_1}$ and $-n\varphi_{C_2}$ about the pulsed spin-lock axes $\hat{C}_{1z'}$ and $\hat{C}_{2z'}$.

Rotating $\mathcal{H}_{CCZQ}$ about *z*, and defining $\Delta\Omega\tau = (\Omega_{C_1} - \Omega_{C_2})(t-n\tau_R)$,

$$\begin{aligned}\omega_{C_1C_2}(T_{11}^{C_1}T_{1-1}^{C_2} + T_{1-1}^{C_1}T_{11}^{C_2}) &\xrightarrow{-\hat{C}_{z1}\Omega_1\tau - \hat{C}_{z2}\Omega_2\tau} \omega_{C_1C_2}(e^{i\Delta\Omega\tau}T_{11}^{C_1}T_{1-1}^{C_2} + e^{-i\Delta\Omega\tau}T_{1-1}^{C_1}T_{11}^{C_2}) \\ &= \sum_{\substack{p=-1\\p\neq 0}}^{1} \omega_{C_1C_2}T_{1p}^{C_1}T_{1-p}^{C_2}e^{ip\Delta\Omega\tau}\end{aligned} \tag{S5}$$

Next, rotating $\mathcal{H}_{CCZQ}$ by $-(\psi_{C_1}+\frac{\pi}{2})$ about $\hat{C}_{1y}$ and $-(\psi_{C_2}+\frac{\pi}{2})$ about $\hat{C}_{2y}$, and collecting terms, we obtain

$$\rightarrow \sum_{q_{C_1}=-1}^{1}\sum_{q_{C_2}=-1}^{1}\sum_{\substack{p=-1\\p\neq 0}}^{1} \begin{matrix}\left(\omega_{C_1C_2}/4\right)\left[\left(1-\delta_{0,q_{C_1}}\right)\left(1-pq_{C_1}\sin(\psi_{C_1})\right) - \delta_{0,q_{C_1}}p\sqrt{2}\cos(\psi_{C_1})\right] \\ \cdot\left[\left(1-\delta_{0,q_{C_2}}\right)\left(1+pq_{C_2}\sin(\psi_{C_2})\right) + \delta_{0,q_{C_2}}p\sqrt{2}\cos(\psi_{C_2})\right]T_{1q_{C_1}}^{C_1}T_{1q_{C_2}}^{C_2}e^{ip\Delta\Omega\tau}\end{matrix} \tag{S6}$$

Finally, rotating $\mathcal{H}_{CCZQ}$ about $\hat{C}_{1z'}$ and $\hat{C}_{2z'}$ by $-n\varphi_{C_1}$ and $-n\varphi_{C_2}$, respectively, and including the MAS time dependence of $\omega_{C_1C_2}$, we obtain:

$$\tilde{\mathcal{H}}_{CCZQ} = \sum_{m=-2}^{2}\sum_{q_{C_1}=-1}^{1}\sum_{q_{C_2}=-1}^{1}\sum_{\substack{p=-1\\p\neq 0}}^{1} \begin{matrix}\left(\omega_{C_1C_2}^{m}/4\right)\left[\left(1-\delta_{0,q_{C_1}}\right)\left(1-pq_{C_1}\sin(\psi_{C_1})\right) - \delta_{0,q_{C_1}}p\sqrt{2}\cos(\psi_{C_1})\right] \\ \cdot\left[\left(1-\delta_{0,q_{C_2}}\right)\left(1+pq_{C_2}\sin(\psi_{C_2})\right) + \delta_{0,q_{C_2}}p\sqrt{2}\cos(\psi_{C_2})\right]T_{1q_{C_1}}^{C_1}T_{1q_{C_2}}^{C_2} \\ \cdot\exp\left(in\left[\varphi_{C_1}q_{C_1} + \varphi_{C_2}q_{C_2}\right]\right)\exp\left(im\omega_R t\right)\exp\left(ip(\Omega_{C_1}-\Omega_{C_2})(t-n\tau_R)\right)\end{matrix} \tag{S7}$$

This interaction-frame Hamiltonian **(S7)** is added to Eq. **(8)** in the main text, when homonuclear transfers are considered for PULSAR.

We now calculate the first-order average Hamiltonian for the flip-flop term. For simplicity, we neglect all prefactors and summations, and concentrate on the time dependences.

$$
\begin{aligned}
\bar{\tilde{\mathcal{H}}}^{(1)}_{CCZQ} &\propto \int_0^{N\tau_R} \exp\left(in\left[\varphi_{C_1}q_{C_1}+\varphi_{C_2}q_{C_2}\right]\right)\exp\left(im\omega_R t\right)\exp\left(ip\Delta\Omega(t-n\tau_R)\right)dt \\
&= \sum_{n=0}^{N-1}\exp\left(in\left[\varphi_{C_1}q_{C_1}+\varphi_{C_2}q_{C_2}\right]\right)\int_{(n-\frac{1}{2})\tau_R}^{(n+\frac{1}{2})\tau_R}\exp\left(im\omega_R t\right)\exp\left(ip\Delta\Omega(t-n\tau_R)\right)dt
\end{aligned}
\tag{S8}
$$

Assuming there is no rotational resonance, $m\omega_R + p\Delta\Omega \neq 0$,

$$
\bar{\tilde{\mathcal{H}}}^{(1)}_{CCZQ} \propto \frac{2(-1)^m \sin(p\Delta\Omega\frac{\tau_R}{2})}{m\omega_R + p\Delta\Omega}\sum_{n=0}^{N-1}\left(e^{i\left[\varphi_{C_1}q_{C_1}+\varphi_{C_2}q_{C_2}\right]}\right)^n \tag{S9}
$$

The summation here is over roots of unity, so is zero unless

$$
\varphi_{C_1}q_{C_1}+\varphi_{C_2}q_{C_2} = 2\pi l \tag{S10}
$$

for some integer $l$. This is the same equality that determines whether the second-order cross terms between terms 1 and 2 are zero, see Eq. **(10)** in the main text, and **SI Section 3**. As such, we conclude that the homonuclear dipolar coupling averages to zero to first order, unless Eq. **(S10)** is fulfilled. We now discuss the possible ways in which it is non-zero, following the same matching conditions analyzed in the main text:

1) If the zero-quantum matching condition is fulfilled, that is, $\varphi_{C_1}-\varphi_{C_2}=2\pi l$ for some integer $l$, then when $q_{C_1}=-q_{C_2}=\pm 1$, Eq. **(10)** and Eq. **(S10)** are satisfied. The 1x2 cross term in Eq. **(9)** produces terms in the second-order average Hamiltonian with spin product operators $T^{H}_{10}T^{C_1}_{11}T^{C_2}_{1-1} \propto H_z C_1^+ C_2^-$ and $T^{H}_{10}T^{C}_{1-1}T^{N}_{11} \propto H_z C_1^- C_2^+$, while the first-order average Hamiltonian of the homonuclear dipolar coupling produces zero-quantum terms $C_1^+C_2^-$ and $C_1^-C_2^+$. Both of these terms should accomplish transfer between the carbons, without a change in sign. The relative sign of the terms determines whether they each are $ZQ_x$ or $ZQ_y$, and thereby whether the terms commute. In either case, numerical simulations indicate that the homonuclear dipolar coupling plays an insignificant role in P-TSAR experiments (**Fig. S3**).

2) If instead the double-quantum matching condition is met, $\varphi_{C_1}+\varphi_{C_2}=2\pi l$ for some integer $l$, then when $q_{C_1}=q_{C_2}=\pm 1$, Eq. **(10)** and Eq. **(S10)** are satisfied. The 1x2 cross term in Eq. **(9)** produces double-quantum product operators $H_z C_1^+ C_2^+$ and $H_z C_1^- C_2^-$, while the first-order average Hamiltonian of the homonuclear dipolar coupling produces double-quantum terms $C_1^+C_2^+$ and $C_1^-C_2^-$. These double-quantum terms cause polarization transfer between the carbons, with a change in sign. Again, whether the homonuclear dipolar coupling and 1x2 cross terms produce commuting operators depends on the relative sign of their terms. But in any case, the homonuclear dipolar coupling plays an insignificant role in P-TSAR (**Fig. S3**).

3) Finally, for any $\varphi_{C_1}$ and $\varphi_{C_2}$, $q_{C_1} = q_{C_2} = 0$, satisfies **(Eq. 10, S10)** trivially. The spin product-operator would be $H_z C_{1z} C_{2z}$ for the 1x2 cross term, and $C_{1z} C_{2z}$ for the first-order homonuclear dipolar coupling. These longitudinal terms commute with both the zero-quantum and double-quantum terms from cases 1 and 2, and will not accomplish any transfer.

In conclusion, the flip-flop term in the homonuclear dipolar coupling averages to zero in first order AHT, unless one of the matching conditions that also allow second-order cross terms between terms 1 and 2 to be satisfied. In any case, the effect of homonuclear dipolar coupling is minimal, based on numerical simulations (**Fig. S3**).

#### 3. Second-order average Hamiltonian of the cross term between terms 1 and 2 of the P-TSAR Hamiltonian

We claim that the second-order cross term between terms 1 and 2 is zero if $e^{i(\varphi_N q_N + \varphi_C q_C)} \neq 1$, i.e. it is non-zero only if $\varphi_N q_N + \varphi_C q_C = 2\pi l$ for some integer $l$. In general, the second-order average Hamiltonian is

$$\bar{\tilde{\mathcal{H}}}^{(2)} = \frac{1}{2iN\tau_R} \int_0^{N\tau_R} dt_2 \int_0^{t_2} dt_1 \left[ \tilde{\mathcal{H}}(t_2), \tilde{\mathcal{H}}(t_1) \right] \quad \text{(S11)}$$

For the cross term between terms 1 and 2,

$$\bar{\tilde{\mathcal{H}}}^{(2)}_{1\times2} = \frac{1}{2iN\tau_R} \int_0^{N\tau_R} dt_2 \int_0^{t_2} dt_1 \left[ \tilde{\mathcal{H}}_1(t_2), \tilde{\mathcal{H}}_2(t_1) \right] - \left[ \tilde{\mathcal{H}}_1(t_1), \tilde{\mathcal{H}}_2(t_2) \right] \quad \text{(S12)}$$

Adding the explicit forms for each interaction frame Hamiltonian from **(8)**, we obtain

$$\bar{\tilde{\mathcal{H}}}^{(2)}_{1\times2} = \sum_{\substack{m_1, q_C, q_{H1} \\ m_2, q_N, q_{H2}}} \left\{ \begin{array}{l} \frac{1}{2iN\tau_R} \omega_{HC}^{m} q_{H1} \left[ q_C \cos(\psi_C) + \delta_{0,q_C} \sqrt{2} \sin(\psi_C) \right] \omega_{HN}^{m} q_{H2} \left[ q_N \cos(\psi_N) + \delta_{0,q_N} \sqrt{2} \sin(\psi_N) \right] \\ T^C_{1q_C} T^N_{1q_N} \left[ T^H_{1q_{H1}}, T^H_{1q_{H2}} \right] \\ \int_0^{N\tau_R} dt_2 \int_0^{t_2} dt_1 \left[ \begin{array}{l} \exp\{ i(n_2 \varphi_C q_C + n_1 \varphi_N q_N) \} \exp\{ i([q_{H1}\omega_{1,H} + m_1\omega_R] t_2 + [q_{H2}\omega_{1,H} + m_2\omega_R] t_1) \} \\ -\exp\{ i(n_2 \varphi_N q_N + n_1 \varphi_C q_C) \} \exp\{ i([q_{H2}\omega_{1,H} + m_2\omega_R] t_2 + [q_{H1}\omega_{1,H} + m_1\omega_R] t_1) \} \end{array} \right] \end{array} \right\}$$

(S13)

Here $n_1 \equiv \lfloor t_1 / \tau_R + 1/2 \rfloor$ and $n_2 \equiv \lfloor t_2 / \tau_R + 1/2 \rfloor$. This $\bar{\tilde{\mathcal{H}}}^{(2)}_{1\times2}$ is repeated in the main text Eq. **(9)**. We note that the commutator is zero unless $q_{H1} = -q_{H2}$. To simplify evaluation, we shift the domain of integration from $t_2 \in [0, N\tau_R]$ to $t_2 \in [\frac{1}{2}\tau_R, (N+\frac{1}{2})\tau_R]$, which will not significantly affect the result, as all terms are $N\tau_R$-periodic.

We neglect, for now, all prefactors before the integrals, and leave the exterior summation implied. To evaluate the integral, we first split into sums over $n_1$ and $n_2$, and replace the argument of the integral with $X$ for clarity:

$$\int_{\frac{1}{2}\tau_R}^{(N+\frac{1}{2})\tau_R} dt_2 \int_{\frac{1}{2}\tau_R}^{t_2} dt_1 X\left(n_2\left(t_2\right),t_2,n_1\left(t_1\right),t_1\right)=\sum_{n_2=1}^{N}\int_{(n_2-\frac{1}{2})\tau_R}^{(n_2+\frac{1}{2})\tau_R} dt_2 \int_{(n_2-\frac{1}{2})\tau_R}^{t_2} dt_1 X\left(n_2,t_2,n_1=n_2,t_1\right) + \sum_{n_2=2}^{N}\sum_{n_1=1}^{n_2-1}\int_{(n_2-\frac{1}{2})\tau_R}^{(n_2+\frac{1}{2})\tau_R} dt_2 \int_{(n_1-\frac{1}{2})\tau_R}^{(n_1+\frac{1}{2})\tau_R} dt_1 X\left(n_2,t_2,n_1,t_1\right) \tag{S14}$$

We will show that if $e^{i(\varphi_N q_N+\varphi_C q_C)}\neq 1$, both of these summations are zero, and hence the entire cross term is zero.

We first evaluate the single sum, in which $n_1=n_2$. To simplify, we substitute $q_{H1}\omega_{1,H}+m_1\omega_R=Q_1$ and $-q_{H1}\omega_{1,H}+m_2\omega_R=Q_2$, and make the $u$-substitutions $T_1=t_1-(n_1-\frac{1}{2})\tau_R$ and $T_2=t_2-(n_2-\frac{1}{2})\tau_R$:

$$\begin{aligned}&\sum_{n_2=1}^{N}\int_{(n_2-\frac{1}{2})\tau_R}^{(n_2+\frac{1}{2})\tau_R} dt_2 \int_{(n_2-\frac{1}{2})\tau_R}^{t_2} dt_1 X\left(n_2,t_2,n_1=n_2,t_1\right)\\ &=\sum_{n_2=1}^{N}\exp\{in_2(\varphi_N q_N+\varphi_C q_C)\}\exp\{i(n_2-\tfrac{1}{2})\omega_R\tau_R(m_1+m_2)\}\int_0^{\tau_R} dT_2\int_0^{T_2} dT_1\begin{bmatrix}\exp(iQ_1T_2+iQ_2T_1)\\ -\exp(iQ_2T_2+iQ_1T_1)\end{bmatrix}\\ &=\int_0^{\tau_R} dT_2\int_0^{T_2} dT_1\begin{bmatrix}\exp(iQ_1T_2+iQ_2T_1)\\ -\exp(iQ_2T_2+iQ_1T_1)\end{bmatrix}(-1)^{m_1+m_2}\sum_{n_2=1}^{N}\left(e^{i(\varphi_N q_N+\varphi_C q_C)}\right)^{n_2}\end{aligned} \tag{S15}$$

The summation over $n_2$ is over roots of unity, as $N\varphi_N$ and $N\varphi_C$ are multiples of $2\pi$. If $e^{i(\varphi_N q_N+\varphi_C q_C)}\neq 1$, this sum is zero, hence the single sum in Eq. **(S14)** is zero.

Consider now the double sum, $n_1\neq n_2$. We concentrate only on the first term within the double integral. Identical arguments hold for the second term. To simplify, we again substitute $q_{H1}\omega_{1,H}+m_1\omega_R=Q_1$ and $-q_{H1}\omega_{1,H}+m_2\omega_R=Q_2$, and make the $u$-substitutions $T_1=t_1-(n_1-\frac{1}{2})\tau_R$ and $T_2=t_2-(n_2-\frac{1}{2})\tau_R$:

$$\begin{aligned}&\sum_{n_2=2}^{N}\sum_{n_1=1}^{n_2-1}\int_{(n_2-\frac{1}{2})\tau_R}^{(n_2+\frac{1}{2})\tau_R} dt_2 \int_{(n_1-\frac{1}{2})\tau_R}^{(n_1+\frac{1}{2})\tau_R} dt_1 X\left(n_2,t_2,n_1,t_1\right)\\ &=\sum_{n_2=2}^{N}\sum_{n_1=1}^{n_2-1}\exp\{in_2(\varphi_C q_C)+in_1(\varphi_N q_N)+i(n_2-\tfrac{1}{2})\tau_R Q_1+i(n_1-\tfrac{1}{2})\tau_R Q_2\}\int_0^{\tau_R}\exp\{iQ_1T_2\}dT_2\int_0^{\tau_R}\exp\{iQ_2T_1\}dT_1\\ &=\int_0^{\tau_R}\exp\{iQ_1T_2\}dT_2\int_0^{\tau_R}\exp\{iQ_2T_1\}dT_1(-1)^{m_1+m_2}\sum_{n_2=2}^{N}\sum_{n_1=1}^{n_2-1}\exp\{in_2A\}\exp\{in_1B\}\end{aligned} \tag{S16}$$

where we have substituted $A = \varphi_C q_C + q_{H1}\omega_{1,H}\tau_R$ and $B = \varphi_N q_N - q_{H1}\omega_{1,H}\tau_R$. Due to the assumptions made to apply AHT, both $e^{iA}$ and $e^{iB}$ are roots of unity and are also not 1, and thus can be written $e^{iA} = e^{2\pi ia/N}$; $e^{iB} = e^{2\pi ib/N}$; $a, b \in \{1, 2, \ldots, N-1\}$. We consider the case $e^{i(A+B)} = e^{i(q_C\varphi_C + q_N\varphi_N)} \neq 1$, or equivalently $a + b \neq N$. We are left to prove the following: For two integers $a, b \in \{1, 2, \ldots, N-1\}$, such that $a + b \neq N$, $\sum_{n_2=2}^{N}\sum_{n_1=1}^{n_2-1}\left(e^{2\pi ia/N}\right)^{n_2}\left(e^{2\pi ib/N}\right)^{n_1} = 0$. We have been unable to find an elementary proof of this claim, so we explicitly calculated all cases for N < 250, and found it to hold. We assume that this trend holds, and note that AHT will be a poor approximation for large *N* in either case. Thus, the double sum is zero if $e^{i(q_C\varphi_C + q_N\varphi_N)} \neq 1$.

In conclusion, the cross term between terms 1 and 2 in the second-order average Hamiltonian is zero unless $e^{i(q_C\varphi_C + q_N\varphi_N)} = 1$. Therefore P-TSAR transfer can occur only when $\varphi_N q_N + \varphi_C q_C = 2\pi l$ for some integer *l*.

**4. Analytical calculation of the finite-pulse spin-lock sequence**

Here we seek analytical formulae for the pulsed spin-lock axis, $\hat{x}'$, and the time-averaged effective pulsed spin-lock frequency, $\omega_{SL} \equiv \varphi / \tau_R$, while accounting for finite pulse effects. In the main text, we considered an isolated spin with chemical shift offset Ω, with a delta-pulse applied at the center of the rotor period with nominal flip angle $\theta$. Now, instead considering a finite pulse, the total rotation due to isotropic chemical shift and the finite rf pulse in one rotor period is

$$R_{x'}(\varphi) = R_z\left(\Omega(\tau_R - \tau_p)/2\right)\cdot R_{\hat{B}_{1,eff}}\left(\theta\cdot\sqrt{1+(\omega_1/\Omega)^2}\right)\cdot R_z\left(\Omega(\tau_R - \tau_p)/2\right) \tag{S17}$$

where $\tau_R$ is the rotor period, $\tau_p$ is the pulse duration, $\omega_1$ is the pulse's nominal nutation frequency, and $\hat{B}_{1,eff} = (\omega_1\hat{x} + \Omega\hat{z})/\sqrt{\omega_1^2 + \Omega^2}$.

We simplify the above equation with the following substitutions: $\hat{B}_{1,eff} = \cos(\zeta)\hat{x} + \sin(\zeta)\hat{z}$, $\delta = \Omega(\tau_R - \tau_p)/2$, and $\theta' = \theta\cdot\sqrt{1+(\omega_1/\Omega)^2}$. Thus $\zeta$ is the angle from the *x*- axis to $\hat{B}_{1,eff}$, $\delta$ is the free evolution time before and after the pulse in each rotor period, and $\theta'$ is the total rotation about $\hat{B}_{1,eff}$ due to the pulse. Our overall rotation is thus

$$R_{x'}(\varphi) = R_z(\delta)\cdot R_{\begin{bmatrix}\cos(\zeta)\\0\\\sin(\zeta)\end{bmatrix}}(\theta')\cdot R_z(\delta) \tag{S18}$$

The rotation matrix about z is:

$$R_z(\delta) = \begin{pmatrix}\cos(\delta) & -\sin(\delta) & 0\\ \sin(\delta) & \cos(\delta) & 0\\ 0 & 0 & 1\end{pmatrix} \tag{S19}$$

While the central rotation matrix in Eq. (S18) is:

$$R_{\begin{bmatrix}\cos(\zeta)\\0\\\sin(\zeta)\end{bmatrix}}(\theta')=\begin{pmatrix}\cos^2(\zeta)+\sin^2(\zeta)\cos(\theta') & -\sin(\zeta)\sin(\theta') & \sin(\zeta)\cos(\zeta)(1-\cos(\theta'))\\ \sin(\zeta)\sin(\theta') & \cos(\theta') & -\cos(\zeta)\sin(\theta')\\ \sin(\zeta)\cos(\zeta)(1-\cos(\theta')) & \cos(\zeta)\sin(\theta') & \sin^2(\zeta)+\cos^2(\zeta)\cos(\theta')\end{pmatrix} \quad \text{(S20)}$$

The overall rotation each rotor period $R_{x'}(\varphi)$ can then be computed as the matrix product in **(S18)**. From $R_{x'}(\varphi)$, we wish to extract the spin-lock axis and angle $\hat{x}'$ and rotation angle per rotor period $\varphi$. For a non-symmetric rotation matrix $A$, the axis of rotation $\vec{u}$ can be extracted:

$$A=\begin{pmatrix}a & b & c\\ d & e & f\\ g & h & i\end{pmatrix}\Rightarrow \vec{u}=\begin{pmatrix}h-f\\ c-g\\ d-b\end{pmatrix} \quad \text{(S21)}$$

The direction of $\vec{u}$ is always such that the matrix is a right-handed rotation described by a rotation angle $\alpha\in(0,\pi)$. To determine $\alpha$, we note $Tr(A)=a+e+i=1+2\cos(\alpha)$. We can thus calculate $\hat{x}'$ and $\varphi$ by replacing $A$ with $R_{x'}(\varphi)$. If $R_{x'}(\varphi)$ is symmetric, the net rotation each rotor period is 0 or 180 degrees: in either case no "spin-lock" direction can be defined, and no TSAR transfer is observed.

To distinguish positive and negative flip-angles, we define $\hat{x}'$ to always have a positive $x$-component. If $\vec{u}$ has a negative $x$-component, $\hat{x}'=-\vec{u}/|\vec{u}|$ and $\varphi=-\alpha$, otherwise $\hat{x}'=\vec{u}/|\vec{u}|$ and $\varphi=\alpha$. The time-averaged pulsed spin-lock frequency can be calculated by dividing $\varphi$ by the rotor period $\tau_R$: $\omega_{SL}\equiv\varphi/\tau_R$. The formulae for $\hat{x}'$ and $\varphi$ are

$$\hat{x}'=sign(\vec{u}\cdot\hat{x})\vec{u}/|\vec{u}|;\quad \varphi=\alpha\, sign(\vec{u}\cdot\hat{x}) \quad \text{(S22)}$$

$$\vec{u}=\begin{pmatrix}2\cos(\delta)\cos(\zeta)\sin(\theta')-2\sin(\delta)\sin(\zeta)\cos(\zeta)(1-\cos(\theta'))]\\ 0\\ 2\sin(\delta)\cos(\delta)[\cos^2(\zeta)+\cos(\theta')(1+\sin^2(\zeta))]+2[\cos^2(\delta)-\sin^2(\delta)]\sin(\zeta)\sin(\theta')\end{pmatrix} \quad \text{(S23)}$$

$$\begin{aligned}\alpha=\arccos(-(1/2)[1+\sin^2(\delta)\cos(\theta')+\cos^2(\zeta)(\sin^2(\delta)-\cos(\theta'))-\sin^2(\zeta)(1-\sin^2(\delta)\cos(\theta'))\\ -\cos^2(\delta)[\cos^2(\zeta)+(1+\sin^2(\zeta))\cos(\theta')]+4\sin(\delta)\cos(\delta)\sin(\zeta)\sin(\theta')])\end{aligned} \quad \text{(S24)}$$

To reiterate, $\vec{u}$ and $\alpha$ are our desired spin lock direction $\hat{x}'$ and rotation angle per rotor period $\varphi$, up to normalization and a sign definition **(S22)**. The MATLAB script in Appendix S3 uses **(S22-24)** to compute the analytical transfer efficiencies in Eq **(14)**.

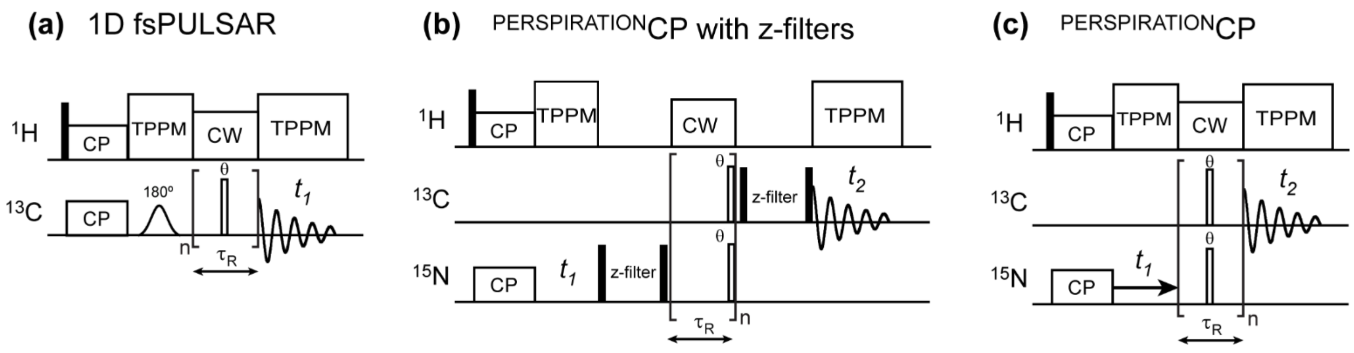


**Figure S1.** Additional pulse sequences used in this study. **(a)** 1D $^{13}$C frequency-selective PULSAR (fsPULSAR) sequence with a long Gaussian 180˚ pulse in an echo following CP to selectively measure Cα magnetization transfer to other carbons. **(b)** The original $^{15}$N-$^{13}$C $^{PERSPIRATION}$CP sequence, where z-filters before and after the pulsed spin-lock period were used to remove unwanted coherences arising from the placement of the pulsed spin-lock pulses at the conclusion of each rotor period. **(c)** The improved $^{15}$N-$^{13}$C $^{PERSPIRATION}$CP sequence where the θ-angle pulsed spin-lock pulses were applied in the middle of each rotor period, without the need for z-filters.

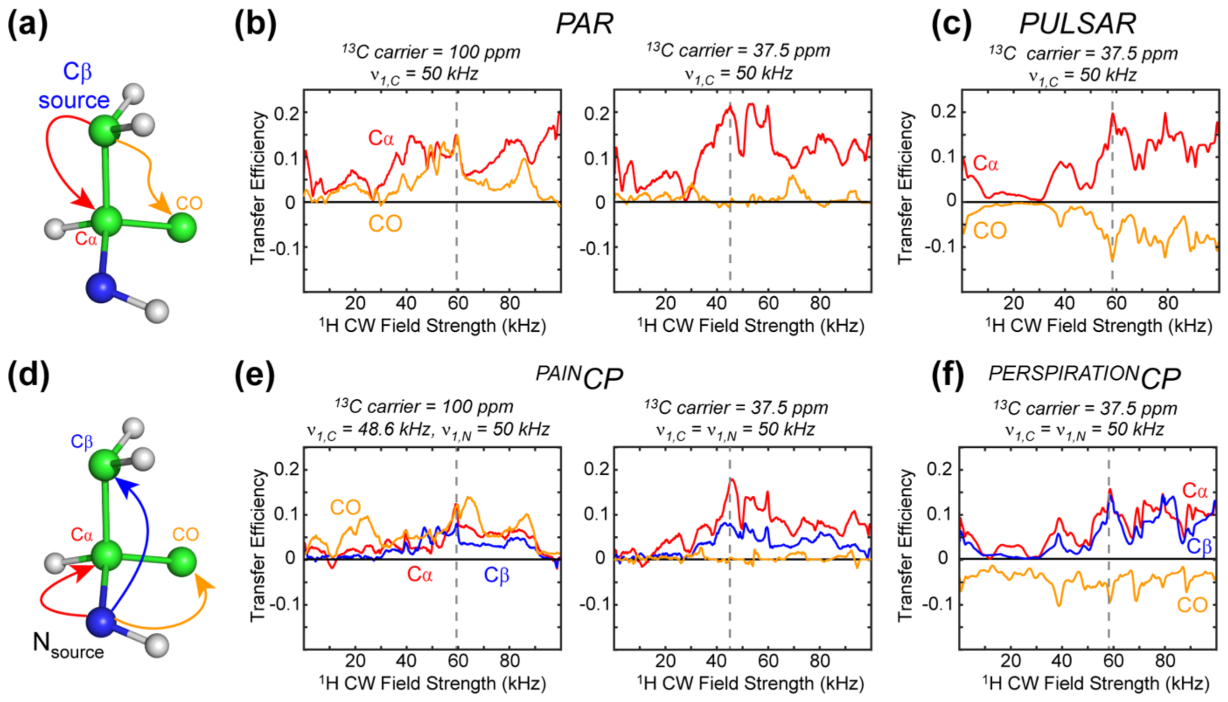


**Figure S2.** Numerical simulations comparing the dependence of P-TSAR and CW-TSAR experiments on $^{1}$H rf field strength. (**a**) Spin system for the PULSAR and PAR simulations. $^{15}$N is omitted from these simulations and the magnetization transfer from $^{13}$Cβ to $^{13}$Cα and $^{13}$CO is tracked. (**b**) PAR transfer efficiency as a function of $^{1}$H rf field. (**c**) PULSAR transfer efficiency as a function of $^{1}$H rf field. (**d**) Spin system for the $^{PERSPIRATION}$CP and $^{PAIN}$CP simulations. An 8-spin system is used (**Table S1**), and magnetization transfer from $^{15}$N to $^{13}$Cα, $^{13}$Cβ, and $^{13}$CO is tracked. (**e**) $^{PAIN}$CP transfer efficiency as a function of $^{1}$H rf field strength. **f)** $^{PERSPIRATION}$CP transfer efficiency as a function of the $^{1}$H rf field strength. All simulations were conducted for a 600 MHz spectrometer using an MAS frequency of 20 kHz.

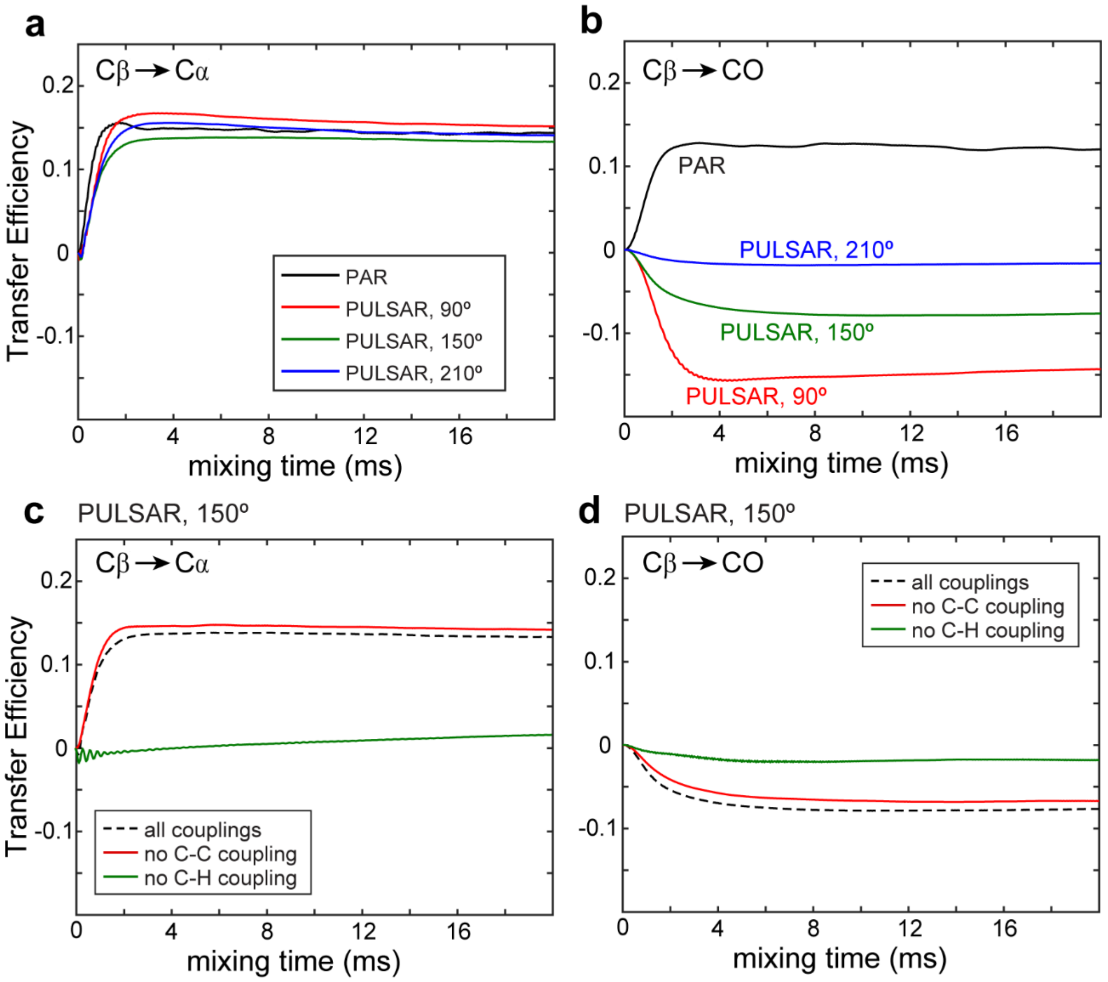


**Figure S3.** Simulated PAR and PULSAR polarization transfer buildup curves. The simulations are for a 600 MHz spectrometer, 20 kHz MAS, 50 kHz $^{13}C$ rf field, 58.5 and 59.5 kHz $^{1}H$ rf fields for PULSAR and PAR, respectively. **(a)** Comparison of PAR and PULSAR buildup curves for Cβ-Cα transfer. **(b)** Comparison of PAR and PULSAR buildup curves for Cβ-CO transfer. **(c, d)** Simulated PULSAR buildup curves for a pulse flip angle of 150˚ for **(c)** Cβ-Cα and **(d)** Cβ-CO transfer under varying conditions of dipolar couplings. These include all couplings are present, when $^{13}C$-$^{13}C$ dipolar couplings are removed, and when $^{13}C$-$^{1}H$ dipolar couplings are removed. Only the $^{13}C$-$^{1}H$ dipolar couplings are necessary for PULSAR polarization transfer.

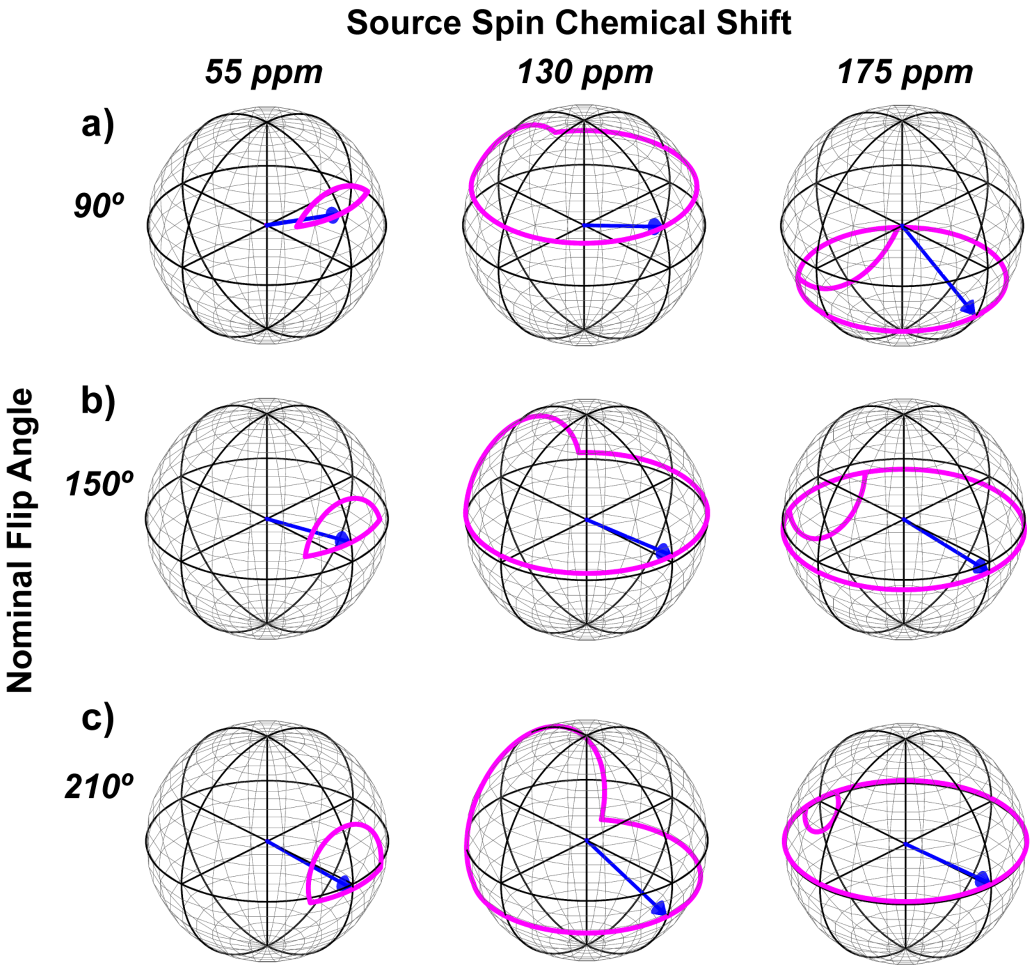


**Figure S4.** Magnetization trajectories along the pulsed spin-lock axis for various pulse flip angles and isotropic chemical shifts of the source spin. **(a)** 90º. **(b)** 150º. **(c)** 210º. The trajectories were calculated using a $^{13}$C rf field of 50 kHz applied on resonance at 37.5 ppm.

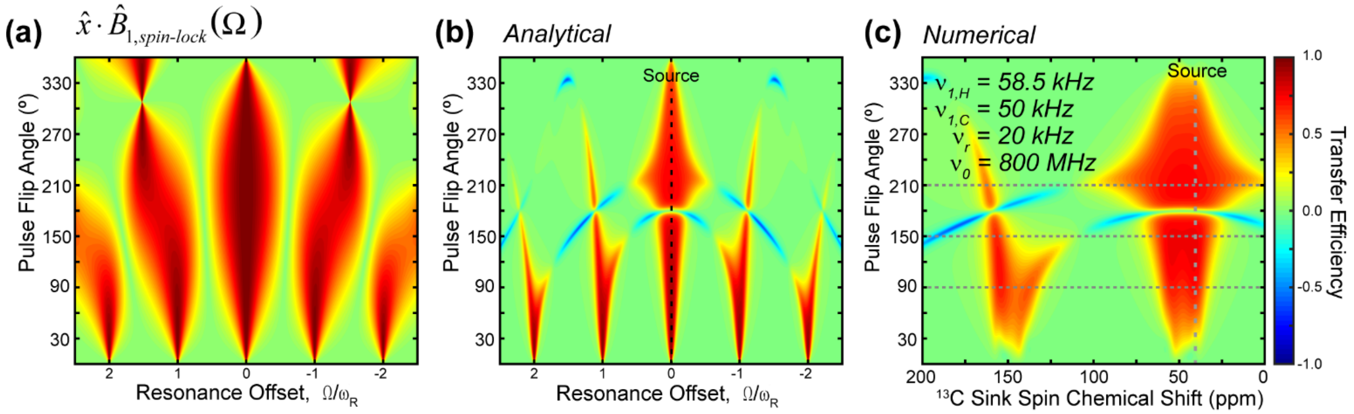


**Figure S5.** Calculated and simulated polarization transfer profile for a P-TSAR sequence where the spin-lock pulse occurs at the end of each rotor period (see pulse sequence in **Fig. S1b**). **(a)** Contour plot showing the "read-in" losses as a function of pulse flip angle and resonance offset. **(b)** Calculated transfer efficiency as a function of chemical shift offset of the sink spin and pulse flip angle. The source spin is on resonance. Positive transfer occurs at the zero-quantum matching condition while negative transfer occurs under the double-quantum condition. Calculations were performed for an 800 MHz spectrometer under 20 kHz MAS with a $^{13}C$ rf field strength of 50 kHz. **(c)** Numerically simulated PULSAR polarization transfer efficiency for when the pulse is placed at the end of each rotor period.

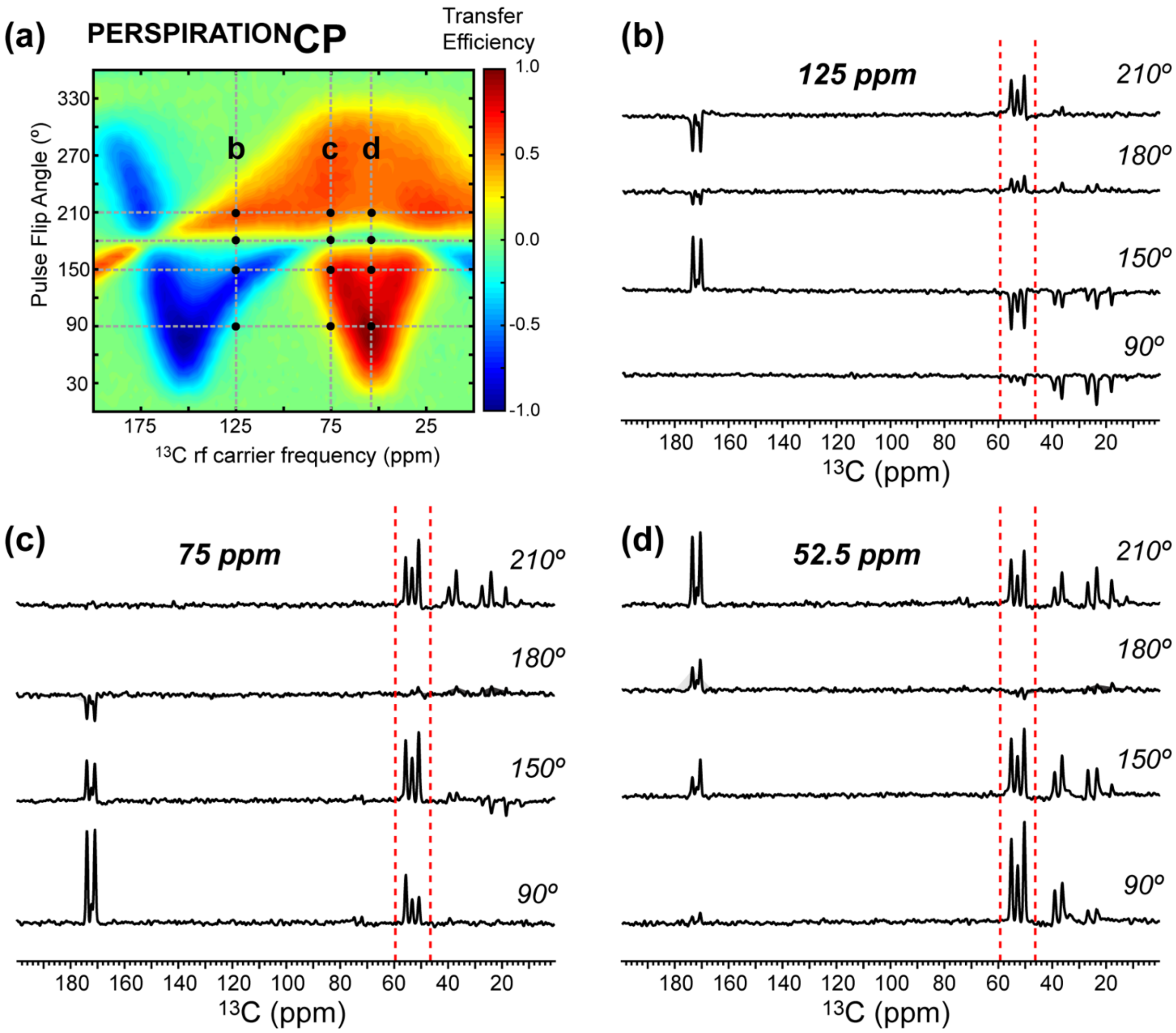


**Figure S6. (a)** 2D heat map of measured $^{PERSPIRATION}$CP integrated intensities for the Cα region of f-MLF. The heat map is constructed from 25 x 80 1D $^{13}C$ spectra after $^{15}N$-$^{13}C$ $^{PERSPIRATION}$CP spectra. The spectra were measured as a function of pulse flip angles at 15˚ increments from 0˚ to 360˚ and $^{13}C$ rf carrier frequencies from 0 to 200 ppm at 2.5 ppm increments. The spectra were measured on an 800 MHz spectrometer under 20 kHz MAS. The $^{1}H$, $^{13}C$ and $^{15}N$ rf field strengths are 57, 47, and 36 kHz, respectively, during the $^{PERSPIRATION}$CP block. **(b)** 1D spectra measured with $^{13}C$ carrier at 125 ppm for 4 different pulse flip angles. **(c)** 1D spectra measured with the $^{13}C$ carrier at 75 ppm. **(d)** 1D spectra measured at $^{13}C$ carrier at 52.5 ppm.

**Table S1**. Atomic coordinates and chemical shift parameters for the 8-spin system used in SpinEvolution simulations of $^{PERSPIRATION}$CP. Simulations for PULSAR omitted the $^{15}N$ nucleus. The atomic coordinates and isotropic chemical shifts are taken from the Met residue of the high-resolution SSNMR structure of f-MLF. Proton CSA was not included in the simulations. CSA parameters for $^{13}C\alpha$, $^{13}C\beta$, and $^{15}N$ were consistent with previous $^{PAIN}$CP simulations [1], while literature reported carbonyl CSA parameters were used for $^{13}CO$ [2]. All simulations were conducted under 20 kHz MAS.

| Nucleus | X, Y, Z (Å) | $\sigma_{iso}$ (ppm) | $\sigma_{11}$, $\sigma_{22}$, $\sigma_{33}$ (ppm) | CSA α, β, γ (°) |
|---|---|---|---|---|
| Cα | 1.246, -0.783, -0.894 | 52 | 77, 39, 39 | 0, 45, 0 |
| Cβ | 0.718, 0.033, 0.320 | 37.5 | 61, 38, 14 | -80, 15, 3 |
| CO | 2.752, -0.881, -0.716 | 172 | 256, 166, 94 | 76, 25, 75 |
| N | 0.678, -2.128. -0.894 | 0 | 98, -42, -55 | 98, 66, 102 |
| $H^N$ | 1.267, -2.967, -0.894 | 0 | - | |
| Hα | 0.953, -0.280, -1.833 | 0 | - | |
| Hβ1 | 1.188, 1.036, 0.300 | 0 | - | |
| Hβ2 | 1.097, -0.432, 1.252 | 0 | - | |

**Appendix S1. SpinEvolution script for varying sink spin chemical shift in $^{PERSPIRATION}$CP**

```
****** The System ***********************************
spectrometer(MHz)  600
spinning_freq(kHz) 20
channels           C13 N15 H1
nuclei             C13 C13 C13 N15 H1 H1 H1 H1
atomic_coords      CNH.cor
cs_isotropic       52 37.5 172 0 0 0 0 0 ppm
csa_parameters     1 25 0 0 45 0 ppm
csa_parameters      2 24 0.92 -80 15 3 ppm
csa_parameters      3 -84 0.85 76 25 75 ppm
csa_parameters      4 98 0.13 98 66 102 ppm
j_coupling         *
quadrupole         *
dip_switchboard    *
csa_switchboard    *
exchange_nuclei    *
bond_len_nuclei    *
bond_ang_nuclei    *
tors_ang_nuclei    *
groups_nuclei      *
******* Pulse Sequence ******************************
CHN 1
timing(usec)      (20.8335 8.333 20.8335)x400
power(kHz)         0 50 0
phase(deg)         0 0 0
freq_offs(kHz)     -6.3 -6.3 -6.3
CHN 2
timing(usec)      (20.8335 8.333 20.8335)
power(kHz)         0 50 0
phase(deg)         0 0 0
freq_offs(kHz)     0 0 0
CHN 3
timing(usec)       (50)
power(kHz)          58.5
phase(deg)          0
freq_offs(kHz)      0
******* Variables ***********************************
scan_par1d cocs/0:0.25:200/
cs_ppm_iso_3 = cocs
******* Options *************************************
rho0               I4x
observables        I1p I2p I3p I4p I5p I6p I7p I8p
EulerAngles        rep168
n_gamma            16
line_broaden(Hz)   *
zerofill           *
FFT_dimensions     *
options            -v1 -re -split64 -clust0
*****************************************************
```

**"CNH.cor" atomic coordinate file used in SpinEvolution simulations of $^{PAIN}$CP and $^{PERSPIRATION}$CP:**

| | | | |
|---|---|---|---|
| 1.246 | -0.783 | -0.894 | C |
| 0.718 | 0.033 | 0.320 | C |
| 2.752 | -0.881 | -0.716 | C |
| 0.678 | -2.128 | -0.894 | N |
| 1.267 | -2.967 | -0.894 | H |
| 0.953 | -0.280 | -1.833 | H |
| 1.188 | 1.036 | 0.300 | H |
| 1.097 | -0.432 | 1.252 | H |

**Appendix S2. SpinEvolution script for simulating 2D PULSAR transfer maps as a function of pulse flip angle and sink spin chemical shift**

```
****** The System ************************************
spectrometer(MHz)  800
spinning_freq(kHz) 20
channels           C13 H1
nuclei             C13 C13 C13 H1 H1 H1 H1
atomic_coords      CNH.cor
cs_isotropic       52 37.5 172  0 0 0 0 ppm
csa_parameters     1 25 0 0 45 0 ppm
csa_parameters      2 24 0.92 -80 15 3 ppm
csa_parameters      3 -84 0.85 76 25 75 ppm
j_coupling         *
quadrupole         *
dip_switchboard    *
csa_switchboard    *
exchange_nuclei    *
bond_len_nuclei    *
bond_ang_nuclei    *
tors_ang_nuclei    *
groups_nuclei      *
******* Pulse Sequence ******************************
CHN 1
timing(usec)      (20.8335 8.333 20.8335)x400
power(kHz)         0 50 0
phase(deg)         0 0 0
freq_offs(kHz)     -9.4 -9.4 -9.4
CHN 2
timing(usec)       (50)
power(kHz)          58.5
phase(deg)          0
freq_offs(kHz)      0
******* Variables ************************************
scan_par1d cocs/0:0.5:200/
scan_par2d p/0:0.3333:19.9999/
pulse_1_1_2=p
pulse_1_1_1=(50-p)/2
pulse_1_1_3=(50-p)/2
cs_ppm_iso_3 = cocs
******* Options **************************************
rho0               I2x
observables        I1p I2p I3p I4p I5p I6p I7p
EulerAngles        rep168
n_gamma            16
line_broaden(Hz)   *
zerofill           *
FFT_dimensions     *
options            -v1 -re -split64 -clust0
*****************************************************
```

**“CNH.cor” atomic coordinate file used in SpinEvolution simulations of PAR and PULSAR:**

```
1.246     -0.783    -0.894    C
0.718     0.033     0.320     C
2.752     -0.881    -0.716    C
1.267     -2.967    -0.894    H
0.953     -0.280    -1.833    H
1.188     1.036     0.300     H
1.097     -0.432    1.252     H
```

**Appendix S3. MATLAB script used for analytical calculations of PULSAR transfer.**

```
%% Efficiency of transferring into or out of the spin-lock
B0_Mhz = 800;
wr_khz = 20;
B1_khz = 50;
beta = 2*pi*B1_khz/wr_khz; % w_effective
carrier = 47.5; % in ppm
thetaList = [0:1:360] ./180.*pi;

%ZQ and DQ transfer width, in units of wr
resWidth = 2/100*2*pi;

ppm = -250:0.5:350; %nominal ppm range
ppmO = ppm - carrier; %ppm offset range
khzO = ppmO * B0_Mhz / 4 / 1000; %

alphaList = khzO./wr_khz.*2.*pi; % offset * tr range, in rad, Iz

x_proj = zeros(length(thetaList), length(alphaList));
z_proj = zeros(length(thetaList), length(alphaList));
etavec = zeros(length(alphaList),1);
phivec = zeros(length(thetaList), length(alphaList));
deltavec = zeros(length(thetaList), length(alphaList));
rotanglevec = zeros(length(thetaList),length(alphaList));
x_projSquared = zeros(length(thetaList), length(alphaList));
chivec = zeros(length(thetaList),length(alphaList));

for idt = 1:length(thetaList)
    theta = thetaList(idt);
    for ida = 1:length(alphaList)
        alpha = alphaList(ida);
        eta = asin(alpha/beta/sqrt((alpha/beta)^2 + 1)); % B1_eff direction up from
x to z (during pulse)
        etavec(ida) = eta;
        phi = theta * sqrt((alpha/beta)^2 + 1); % flip angle during pulse
        phivec(idt,ida) = phi;
        delta = alpha/2 * (1 - theta/beta); % z-rotation during delay
        deltavec(idt,ida) = delta;
        xvec = [2*cos(delta)*cos(eta)*sin(phi) - 2*sin(delta)*sin(eta)*cos(eta)*(1-
cos(phi)),
                0,
                2*sin(delta)*cos(delta)*(cos(eta)^2+cos(phi)*(1+sin(eta)^2)) + ...
                2*(cos(delta)^2-sin(delta)^2)*sin(eta)*sin(phi)]; % unnormalized
spin-lock direction

        rotanglevec(idt,ida) = asin(sqrt(xvec'*xvec)/2);

        chivec(idt,ida) = acos(-
1/2*(1+cos(phi)*sin(delta)^2+cos(eta)^2*(sin(delta)^2-cos(phi)) ...
            -sin(eta)^2 ...
            +cos(phi)*sin(delta)^2*sin(eta)^2-
cos(delta)^2*(cos(eta)^2+cos(phi)*(1+sin(eta)^2)) ...
            +4*cos(delta)*sin(delta)*sin(eta)*sin(phi))) * sign(xvec(1)) +
(sign(xvec(1))==-1)*2*pi; % rotation about spin-lock direction

        xvec = xvec/norm(xvec);
        x_proj(idt,ida) = xvec(1);
        z_proj(idt,ida) = xvec(3);
        x_projSquared(idt,ida) = xvec(1)*xvec(1)*sign(xvec(1));
    end
```

```
end

pulspinlock = zeros(length(thetaList), length(alphaList));
transferonres = zeros(length(thetaList), length(alphaList));
transferCb = zeros(length(thetaList), length(alphaList));

for idt = 1:length(thetaList)
    pulspinlock(idt,:) = x_proj(idt,:) * x_proj(idt, 1+250*2+47.5*2).*...

(x_proj(idt,:).*x_proj(idt,1+250*2+47.5*2)+z_proj(idt,:).*z_proj(idt,1+250*2+47.5*2
));

    transferonres(idt,:) = x_proj(idt,:)*x_proj(idt,1+250*2+47.5*2).* ...

(x_proj(idt,:).*x_proj(idt,1+250*2+47.5*2)+z_proj(idt,:).*z_proj(idt,1+250*2+47.5*2
)).*...
        (1-thetaList(idt)/beta).* ...
        (( resWidth^2./(( chivec(idt,:) - chivec(idt,1+250*2+47.5*2)).^2 +
resWidth^2)) -...
        ( resWidth^2./(( chivec(idt,:) + chivec(idt,1+250*2+47.5*2)-2*pi).^2 +
resWidth^2)) -...
        ( resWidth^2./(( chivec(idt,:) + chivec(idt,1+250*2+47.5*2)-4*pi).^2 +
resWidth^2)));

    transferCb(idt,:) = 1/2*x_proj(idt,:).*(1-thetaList(idt)/beta).*...
        (x_proj(idt,1+250*2+37.5*2).* ...

(x_proj(idt,:).*x_proj(idt,1+250*2+37.5*2)+z_proj(idt,:).*z_proj(idt,1+250*2+37.5*2
)).*...
        ( resWidth^2./(( chivec(idt,:) - chivec(idt,1+250*2+37.5*2)).^2 +
resWidth^2) - ...
        ( resWidth^2./(( chivec(idt,:) + chivec(idt,1+250*2+37.5*2)-2*pi).^2 +
resWidth^2)) -...
        ( resWidth^2./(( chivec(idt,:) + chivec(idt,1+250*2+37.5*2)-4*pi).^2 +
resWidth^2))) ...
        ...
        + x_proj(idt,1+250*2+52*2).* ...

(x_proj(idt,:).*x_proj(idt,1+250*2+52*2)+z_proj(idt,:).*z_proj(idt,1+250*2+52*2)).*
...
        ( resWidth^2./(( chivec(idt,:) - chivec(idt,1+250*2+52*2)).^2 + resWidth^2)
- ...
        ( resWidth^2./(( chivec(idt,:) + chivec(idt,1+250*2+52*2)-2*pi).^2 +
resWidth^2)) - ...
        ( resWidth^2./(( chivec(idt,:) + chivec(idt,1+250*2+52*2)-4*pi).^2 +
resWidth^2))));
end

figure (1)
contourf(ppmO./100, thetaList./pi.*180, pulspinlock, 200, 'edgecolor','none')
colormap('jet')
axis([-2.5 2.5 0 360])
caxis([-1 1])
title('Pulsed Spin-Lock Max Intensity')
colorbar
xlabel('Sink Spin CS (ppm)')
ylabel('Nominal Flip Angle (deg)')

figure (2)
contourf(ppmO./100, thetaList./pi.*180, transferonres, 200, 'edgecolor','none')
```

```
colormap('jet')
axis([-2.5 2.5 0 360])
caxis([-1 1])
title('Transfer Efficiency From Spin at Carrier Position')
xlabel('Sink Spin CS (ppm)')
ylabel('Nominal Flip Angle (deg)')

figure (3)
contourf(ppm, thetaList./pi.*180, transferCb, 200, 'edgecolor','none')
colormap('jet')
axis([0 200 0 360])
caxis([-1 1])
title('Transfer Efficiency From Spin at 37.5 ppm, Carrier at 47.5 ppm')
colorbar
xlabel('Sink Spin CS (ppm)')
ylabel('Nominal Flip Angle (deg)')

figure (4)
for idt = [91, 151, 221]
    plot(ppm,chivec(idt,:)./pi.*180, 'DisplayName', string(thetaList(idt)/pi*180))
    hold on
end
axis([0 200 0 360])
title('Rotation About Spin Lock Axis')
legend
xlabel('CS (ppm)')
ylabel('Rotation (deg)')

figure (5)
for idt = [91, 151, 181, 211]
    plot(ppm,x_proj(idt,:), 'DisplayName', string(thetaList(idt)/pi*180))
    hold on
end
axis([0 200 -1 1])
title('X projection of Pulsed Spin Lock')
legend
xlabel('CS (ppm)')
ylabel('X projection of Pulsed Spin Lock')
```

### Appendix S4. Bruker pulse program for 2D $^{15}$N-$^{13}$C $^{PERSPIRATION}$CP.

```
;p3:        1H excitation pulse
;p15:       CP contact pulse
;p16:       15N PERSP pulse
;p17:       13C PERSP pulse
;p31:       TPPM pulse
;pl2:       1H excitation power
;pl3:       15N CP power
;pl4:       1H CP power
;pl1:       13C PERSP power
;pl5:       15N PERSP power
;pl6:       1H PERSP CW power
;pl12:      1H dec. power during t1 and t2
;cpdprg2:   dec. for t1 evl and t2 acq.
;spnam0:    15N ramp for 1H-15N CP
;d0:        t1 evolution time, initial value set to 0.5 us
;d1:        recycle delay
;d5:        PERSP mixing time
;cnst31:    MAS frequency in Hz
;l5:        PERSP rotor cycles - 2

   "d5 = (l5+2)*1s/cnst31"
   "p12=0.5s/cnst31-p16/2"
   "p13=p12 - 0.3u"  ;p13 is initial read into spin lock
   "p14=p12 - 1u - 5u"  ;p14 is final read out of spin lock
   "p30=p31-0.4"

  "in0=inf1"
  "in10=inf2"

"acqt0=0"                ;for SL; after 90 pulse, acqt=-p1*2/pi

   dccorr
1 ze
  d5
2 d1 do:f2
  10u reset:f1 reset:f2 reset:f3

1m
if "p15<10.1m" goto Passp15
print "contact time exceeds 10msec limit!"
goto HaltAcqu
Passp15, 1m

1m
if "aq<50.1m" goto Passaq
print "acquisition time exceeds 50m limit!"
goto HaltAcqu
Passaq, 1m


  10u pl1:f1
```

```
  10u pl2:f2
  10u pl3:f3
  6u setnmr4|31 \n 4u  setnmr4^31
  10u

3 p3:f2 ph1
  0.3u pl4:f2

;  (p15 ph2):f3 (p15:spf0 ph10):f2
  (p15:spf0 ph2):f3 (p15 ph10):f2
4 1u pl12:f2 cpds2:f2

  d0
  0.3u do:f2

;  ----------------  PERSP block ----------------
; -------- 1st and last cycles are written explicitly so they can be adjusted for phasing ------

  (p13 ph17 pl6):f2
  (center
    (p16 ph7 pl5):f3
    (p16 ph17 pl6):f2
    (p17 ph6 pl1):f1
  )
  (p12 ph17 pl6):f2


5 (p12 ph17 pl6):f2
  (center
    (p16 ph7 pl5):f3
    (p16 ph17 pl6):f2
    (p17 ph6 pl1):f1
  )
  (p12 ph17 pl6):f2
lo to 5 times l5

  (p12 ph17 pl6):f2
  (center
    (p16 ph7 pl5):f3
    (p16 ph17 pl6):f2
    (p17 ph6 pl1):f1
  )
  (p14 ph17 pl6):f2

; ------------ detection ------------
1u pl12:f2 cpds2:f2
 go=2 ph31
 1m do:f2
 10m mc #0 to 2
 F1PH(calph(ph2, +90), caldel(d0, +in0))
 1m do:f2

HaltAcqu, 1m
exit
```

```
ph0= 0
ph1= 1 3                    ;1H 90
ph10= 0                     ;1H CP; M=0 2
ph2= 0                      ;15N CP, M = 0 2
ph6 = 0 0 0 0 2 2 2 2
ph7= 0 0 2 2
ph17 = 0
ph31= 0 2 2 0 2 0 0 2
```

### Appendix S5. Bruker pulse program for 2D $^{13}C$-$^{13}C$ PULSAR

```
;2D PULSAR experiment

;p1:      X Channel hard 90
;p16:     PULSAR contact pulses
;p3:      H1 excitation pulse length
;p15:     CP contact time at pl1 (f1) and pl4 (f2)
;p16:     13C PULSAR pulse length
;pl1:     carbon cp power
;pl2:     H1 excitation pulse power
;pl4:     proton cp power
;pl12:    1H decoupling power
;pl14:    PULSAR power on 1H, ~ 2.9*w_r
;pl15:    13C PULSAR ower
;l5 :     number of rotor cycles for Pulsar - 2
;cnst31:  spinning speed in Hz
;cnst15:  13C carrier (ppm) for PULSAR
;ns :     8*n
;MC2:     States-TPPI
;ND0:     1
;d5:      Pulsar mixing time

  "p12=0.5s/cnst31-p16/2"
  "d5 = (l5+2)*1s/cnst31"
  "p2=2*p1"
  "p30=p31-0.4u"
  "d21 = 0.5s/cnst31-2u-6u"
  "p13 = p12 - 3u"
  "p14 = p12 - 2u - 5u"


   dccorr

1 ze
  d5
2 d1 do:f2
  10u reset:f1 reset:f2
  10u pl1:f1
  10u pl2:f2
  2u:f1 ph2
  2u:f2 ph1
  6u setnmr4|31 \n 4u  setnmr4^31

  p3:f2 ph1
  (p15:spf0 ph2):f1 (p15 pl4 ph10):f2
  1u pl12:f2  cpds2:f2
  d0
  (1u fq=cnst15(bf ppm)):f1
  1u do:f2

;-----------------------------PULSAR mixing----------------------
  (p13 pl14 ph16):f2
  (p16 pl14 ph16):f2 (p16 pl15 ph6):f1
```

```
  (p12 pl14 ph16):f2

5 (p12 pl14 ph16):f2
  (p16 pl14 ph16):f2 (p16 pl15 ph6):f1
  (p12 pl14 ph16):f2
 lo to 5 times l5

  (p12 pl14 ph16):f2
  (p16 pl14 ph16):f2 (p16 pl15 ph6):f1
  (p14 pl14 ph16):f2

;-------------------------  detection ----------------------

  1u cpds2:f2 pl12:f2
  (1u fq=0):f1

  go=2 ph31
  1m do:f2

 10m mc #0 to 2
 F1PH(calph(ph2, +90), caldel(d0, +in0))
 1m do:f2

HaltAcqu, 1m
exit

ph1 = 1 3
ph10= 0
ph2 = 0
ph6  = 0 0 2 2
ph16= 0 0 0 0 2 2 2 2
ph31= 0 2
```

**References**

1. Lewandowski, J. R.; De Paëpe, G.; Griffin, R. G., Proton Assisted Insensitive Nuclei Cross Polarization. *J. Am. Chem. Soc.* **2007,** *129* (4), 728-729.

2. Federico, C.; Karine, L.; Philippe, P.; Geoffrey, B., Determination of Chemical Shift Anisotropy Tensors of Carbonyl Nuclei in Proteins through Cross-Correlated Relaxation in NMR. *Chem. Phys. Chem.* **2004,** *5* (6), 807-814.
